# Visinity: Visual Spatial Neighborhood Analysis for Multiplexed Tissue Imaging Data

**DOI:** 10.1101/2022.05.09.490039

**Authors:** Simon Warchol, Robert Krueger, Ajit Johnson Nirmal, Giorgio Gaglia, Jared Jessup, Cecily C. Ritch, John Hoffer, Jeremy Muhlich, Megan L. Burger, Tyler Jacks, Sandro Santagata, Peter K. Sorger, Hanspeter Pfister

## Abstract

New highly-multiplexed imaging technologies have enabled the study of tissues in unprecedented detail. These methods are increasingly being applied to understand how cancer cells and immune response change during tumor development, progression, and metastasis, as well as following treatment. Yet, existing analysis approaches focus on investigating small tissue samples on a per-cell basis, not taking into account the spatial proximity of cells, which indicates cell-cell interaction and specific biological processes in the larger cancer microenvironment. We present Visinity, a scalable visual analytics system to analyze cell interaction patterns across cohorts of whole-slide multiplexed tissue images. Our approach is based on a fast regional neighborhood computation, leveraging unsupervised learning to quantify, compare, and group cells by their surrounding cellular neighborhood. These neighborhoods can be visually analyzed in an exploratory and confirmatory workflow. Users can explore spatial patterns present across tissues through a scalable image viewer and coordinated views highlighting the neighborhood composition and spatial arrangements of cells. To verify or refine existing hypotheses, users can query for specific patterns to determine their presence and statistical significance. Findings can be interactively annotated, ranked, and compared in the form of small multiples. In two case studies with biomedical experts, we demonstrate that Visinity can identify common biological processes within a human tonsil and uncover novel white-blood cell networks and immune-tumor interactions.

## 1 Introduction

Tobler’s first law of geography, *“Everything is related to everything else, but near things are more related than distant things”* [86] emphasizes the importance of spatial proximity. This is not limited to geographical phenomena; it is also applicable to many biological systems [61]. Biological tissues comprise numerous cell types that function together in multi-cellular units that are crucial in development, physiology, and disease. In cancer, the interactions between tumor cells and immune cells are of particular interest as these contacts dictate whether tumor growth is controlled or proceeds unrestrained [23]. Recent tissue imaging methods permit the identification and quantification of tumor and immune cell types within cancer tissue. Important functional interactions between cells can be inferred by identifying cells that are next to each other. In addition, higher-order arrangements of cells that may represent a structural or functional component of tissue can be quantified by determining which cells tend to neighbor each other (spatial neighborhood analysis). These ‘recurrent neighborhoods’ can assemble to compose more extensive spatial patterns.

Detecting such patterns poses substantial challenges; recent research found that spatial patterns must be investigated across large regions of tissues to yield biologically and statistically meaningful results [50], necessitating methods capable of spatial analysis at large scales.

Experts in the fields of pathology, cancer biology, and systems pharmacology have thus acquired whole-slide images from tissue sections using fluorescence microscopy techniques, such as CyCIF [48], with an overall size of up to 60k × 60k pixels. This results in highly multiplexed tissue images, often larger than 100GB in size. Moreover, a single experiment can involve imaging ten or more specimens within a larger cohort. Identifying spatial neighborhood patterns in such data requires scalable computational methods. However, such techniques alone cannot fully replace the human mind; experts have expansive domain knowledge of cell and tissue morphology formed through years of visually investigating tissues. There is thus a need to facilitate visual human-in-the-loop data exploration, permitting these experts to guide pattern identification and verification. Yet existing visual approaches [77, 84, 85] for spatial neighborhood analysis, by design, only scale to single images representing small regions of tissue and are limited in their interactive capabilities.

We addressed these challenges as a team of visualization researchers, pathologists, and cell biologists via a process of goal specification, iterative design, and tool deployment in a biomedical research laboratory. We make the following contributions: **(1)** A domain-specific human-in-the-loop workflow to visually analyze, extract, and summarize spatial interaction patterns within and across datasets. This workflow enables both exploratory and confirmatory analysis through semi-automatic pattern detection and visual querying. Identified patterns can be annotated with information about their biological context, compared, and saved for continued study. **(2)** A scalable and flexible computational pipeline to quantify cellular neighborhoods and their spatial arrangements into larger spatial microenvironments (patterns). This pipeline quantifies the spatial neighborhood of each cell as a vector of surrounding cell types in a defined query range. We group similar neighborhoods and verify their significance through permutation testing, allowing for the identification of meaningful spatial patterns within and across tissues. (3) A scalable visual analytics system named *Visinity* to interactively analyze the computed neighborhood patterns in and across the large whole-slide tissue image data. Visinity consists of a web-based multiplex image viewer with different rendering modes and superimposed neighborhood encodings. Image exploration is linked to projections and parallel coordinates highlighting frequent neighborhoods and their composition. Small multiple arrangements of these views summarize findings and allow for side-by-side comparison.

We evaluate the applicability of our approach in two hands-on case studies with biomedical experts. We first demonstrate that our system can detect well-established spatial patterns of immune cells in a healthy human tonsil specimen. Second, we analyze a cohort of specimens from a genetically engineered mouse model of lung cancer, revealing immune cell interactions that are an area of cutting-edge research in oncology. We report on user feedback on Visinity’s functionalities and demonstrate the tool’s computational scalability.

## 2 Related Work

### 2.1 Visual Spatial Analysis of Biomedical Imaging Data

A wide variety of bioimaging data viewers (e.g., OMERO Pathviewer [35], ViV [53]), Napari [82], Cytomine [75], Minerva [70]) and visual analysis tools (e.g., ParaGlyder [57], Vitessce [19], Facetto [40], Scope2Screen [32]) are used to study multiplexed tissue images and derived feature data. Visualization and analysis methods for older spatially resolved modalities, by contrast, often operate directly on the pixel data, though not at the single-cell resolution [6, 7, 13, 17]. Generally, the aforementioned tools focus on visual exploration and cell-type identification and are not intended to analyze interactions between cell types and the larger spatial neighborhoods that tissue microenvironments are composed of.

A small subset of tools go beyond single-cell analysis to investigate neighborhood patterns. CytoMAP [85] is a computational toolbox designed to analyze spatial patterns in highly-multiplexed imaging data. Similar to our approach, it uses radial queries to compute local neighborhoods and visualizes their composition and arrangements. However, the static plots that CytoMAP provides do not allow for interactive exploration and search. ImaCytE [84], HistoCat [77], and Halo [2] offer interactive spatial analysis capabilities through linked views. These approaches visualize cell-cell interaction as matrices, superimposed links in the image space (Halo), interaction networks (HistoCAT), and as aggregated glyph-based representations of frequent neighborhoods (ImaCytE). While the proposed visual encodings were evaluated as effective for exploring cell interactions, their confirmatory analysis capabilities are limited. Visinity, by contrast, offers various visual querying capabilities to search for specific interactions.

Most existing systems do not scale to the large datasets our users work with. ImaCytE supports computation and rendering of tens of thousands of cells, whereas whole-slide tissue images often contain upwards of a million cells. Visinity enables this through scalable multi-resolution WebGL rendering, spatial indexing, and algorithms that operate at interactive rates. Additionally, like CytoMAP, most approaches offer isolated analysis of one tissue at a time, whereas Visinity enables users to analyze and compare across specimens. Somarakis et al. [83] support such cohort analysis of pairwise cell-cell interactions through explicit visual encodings (raincloud plots and heatmaps), but data sizes are limited to 10^5^ cells per dataset (Visinity scales to 10^7^ cells). Finally, to test the statistical significance of identified spatial patterns, HistoCAT and ImaCYtE rely on permutation testing. We extend these methods with efficient parallelization, precomputation, and visualization to make them scalable, understandable, and interactively adjustable.

### 2.2 Visualization of Spatial Interaction

Our approach also draws more broadly on work visualizing spatial interactions outside of the biomedical domain, such as movement and communication between geographic areas. A straightforward approach is to display such interactions in their spatial dimensions, e.g., on top of a map or image. Advantages of this approach are the familiarity of reading maps as well as emphasizing the spatial auto-correlation in the interaction data. ArcGIS [1], among other geographical information systems, offers statistical methods and visual encodings [3, 79] to compute and display spatial dependencies, involving spatial correlation, clustering, and alignment of spatial objects (shape, center, orientation). Results are usually superimposed on the map. To show spatial interaction, flow maps [97] are a common practice. Varying opacity [94], spatial aggregation [68, 90], and edge bundling [68] are common methods to resolve clutter in these views. However, while flow maps are well suited for tracking interactions across large distances, cell interactions in tissue usually form a more planar graph with local connectedness. Glyph overlays [74] can indicate a direction (i.e., tensor fields) and additional features without obscuring the underlying data. In our data, interaction is not explicitly defined but indicated through spatial proximity. We thus decided not to emphasize interaction by visual edges or glyphs. Instead, we use minimalistic color-coding to highlight cell types in user-based selections and contours (concave-hulls) to emphasize the unity of detected neighborhood clusters while keeping the underlying image data visible. Interaction patterns have also been displayed in abstract (non-spatial) views where they can be visualized in aggregation, such as node-link diagrams [8] and matrices [97]. Other systems [25, 42], similar to Visinity, apply a combination of coordinated spatial and abstract views [73], offering different perspectives on the data.

Visinity also draws on visual querying to search spatial interaction patterns. PEAX [44] introduces visual querying for pattern search in sequential data where users query by example and interactively train a classifier to find similar patterns. Krueger et al. [69] propose a visual interface to sketch, query, display, and refine spatial interactions between moving objects. We adapt and task-tailor this workflow; search can be triggered by selecting existing patterns in the tissue image or by explicitly sketching a spatial neighborhood composition.

### 2.3 Computational Spatial Analysis Methods

Relevant spatial analysis methods can be categorized into: (A) statistical methods measuring spatial distribution/ correlation of single or pairwise features, (B) approaches detecting higher-order feature interaction such as topics and motifs from tabular data, and (C) image-based approaches to find reoccurring spatial features.

#### A) Spatial (Auto-)Correlation Methods

Moran’s I [56] and Geary’s C [18] are commonly used methods to quantify spatial autocorrelation, which determines how a variable is distributed spatially. Such correlation can be computed for every data feature (e.g., gene/protein expression) in isolation to describe their spatial organization and has been used to identify biological relationships in imaged tissue [13]. Ripley’s K function [72], among others, extends this by computing random, dispersed, or clustered distribution patterns at varying scale, for one (univariate) or between two features (bivariate).

#### B) Higher-order Interactions between Multiple Features

To identify higher-order patterns, the spatial relatedness of objects can also be modeled as a network. Ribeiro et al. summarize the field of sub-graph counting and motif discovery [71]. However, the number of cells (nodes in the graph) in combination with a variety of cell types (node attributes) renders current motif discovery algorithms computationally infeasible for interactive setups, especially when considering interaction at multiple scales. The complexity of most algorithms grows exponentially with motif sizes [93]. Other approaches compute groups (topics or clusters) based on probabilities and distances. Zhu et al. [98] use a Hidden-Markov random field to model spatial dependency of gene expression. Spatial-LDA [91] is a probabilistic topic modeling approach, which is also applied in the biomedical field [58]. CytoMAP [85] and stLearn [67] rely on distance-based clustering; they extract per-cell features from histology images, collect neighborhood information for each cell (represented as vectors), and cluster these vectors to reveal patterns. We formalize and combine these concepts into a computational neighborhood quantification pipeline. We improve existing methods by inverse distance weighting, adding similarity search, and creating a scalable implementation; neighborhood size can be changed on the fly to analyze spatial patterns at different scales.

#### C) Image-based Approaches

Other methods directly operate on the image. Common deep learning approaches [76] include representation learning for comparing and finding similar image regions [14, 95] and convolutional neural networks [21] for object classification and localization. However, supervised approaches are difficult in a biomedical context because of a lack of labeled data and their explainability. Overcoming segmentation by operating directly on pixel data [62, 64, 89] renders interpretation of interactions between captured patch-like structures more challenging. This aligns with our experts’ practice of using cells as elemental and meaningful biological building blocks. By building our approach on single-cell data derived from multiplexed images, we were able to develop a more scalable and quantifiable approach.

## 3 Whole-slide Multiplexed Tissue Imaging Data

The data created by our biomedical experts consists of multiplexed images of tissue generated by iterative staining with antibodies that recognize specific proteins followed by imaging with a high-resolution optical microscope in successive cycles. Our collaborators commonly use cyclic immunofluorescence (CyCIF) [49] to generate these data, though other imaging technologies [20, 22] are capable of producing similar image data. Individual cells in the image are then segmented. Based on the relative protein expression levels present within each cell, most cells can be assigned to a specific cell type [45, 60]. This process, therefore, yields the following data for a single specimen (Fig. 2): **(a)** *a multi-channel tissue image*, where each channel corresponds to a different ‘marker’ (often proteins are recognized by an antibody), **(b)** *a cell segmentation mask* that conveys the location of each individual cell in the image space, and **(c)** *single-cell data*: a feature table that includes position and cell type for each cell. **(d)** Specimens can be parts of greater ‘cohorts’ and are investigated in conjunction to one another. These data are sizeable; single slides range from 2 to 6 cm^2^ in size and contain up to 10^7^ cells. The tissue image, given the number of channels, contains up to 10^9^ pixels, resulting in an image file as large as 200GB. All analysis in this paper uses the OME standard for microscopy image data [46] and is generated with the MCMICRO [78] image processing pipeline.

**Fig. 1:**
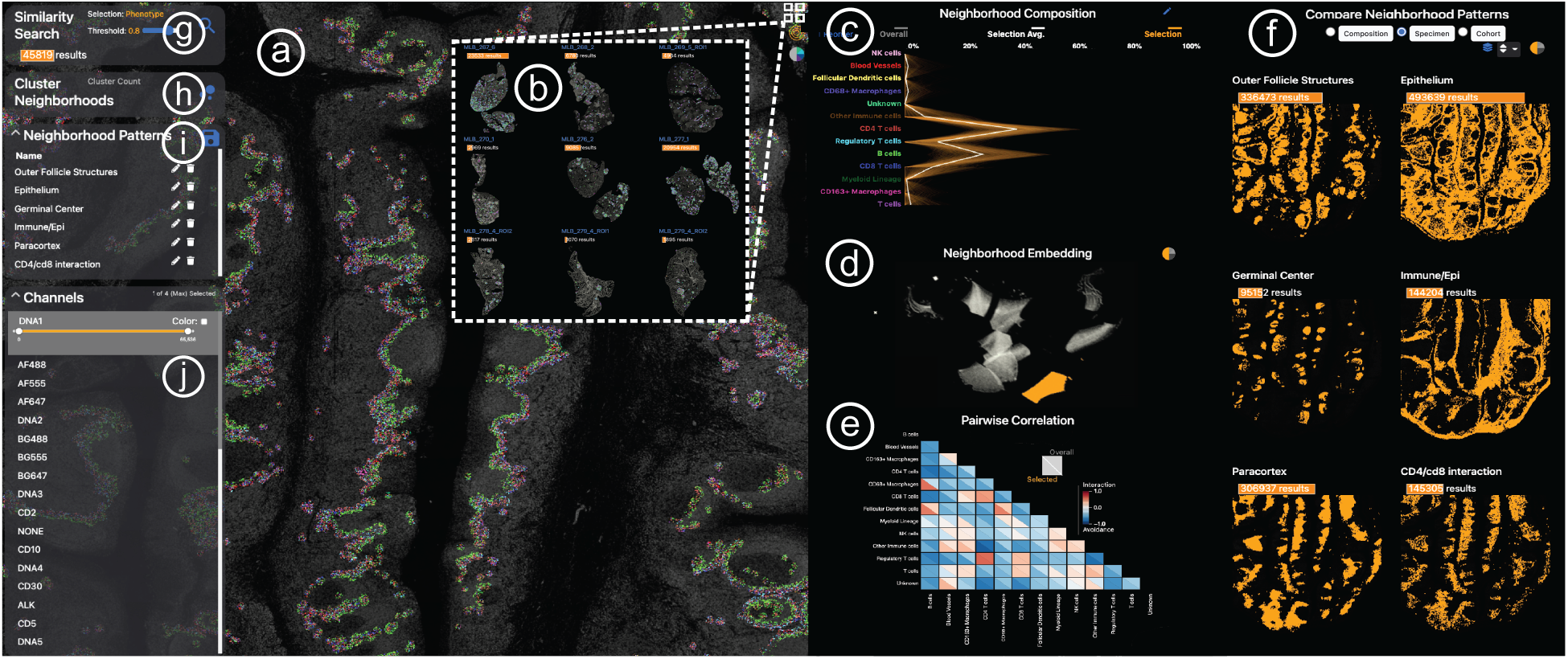
Visinity interface. a) Image viewer: multiplex whole-slide tissue images highlighting spatial cell arrangement; b) Cohort view: search, apply, compare spatial patterns across different specimens; c) Neighborhood composition view: visualizes cell types that make up cell neighborhoods; d) UMAP embedding view: encodes cells with similar neighborhood as dots close to each other; e) Correlation matrix: pairwise interactions between cells; f) Comparison & summary view: different small multiple encodings of extracted patterns; g) Neighborhood search: finds cells with similar neighborhood; h) Interactive clustering: automated detection of neighborhood patterns; i) Annotation panel: save and name patterns; j) Channel selection: color and combine image channels.

**Fig. 2:**
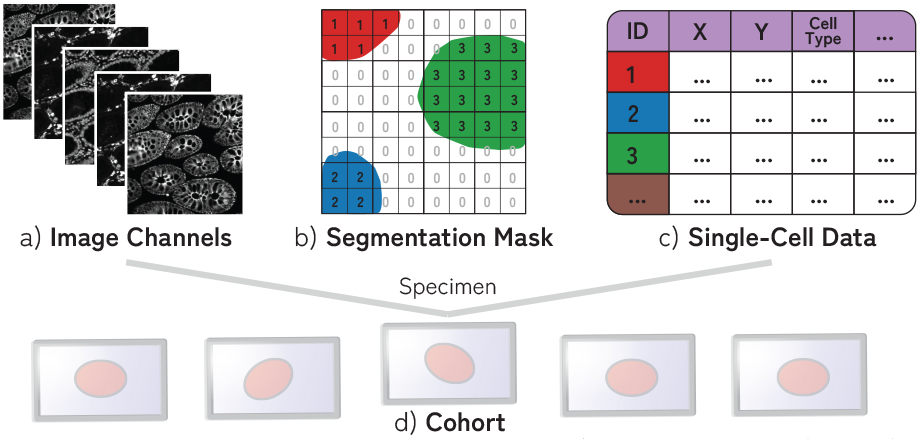
A specimen consists of (a) multi-channel image data, (b) segmentation mask of cells (often > 10^6^ cells), and (c) single-cell data containing information about the position, cell type, and marker intensity values for each cell. (d) Specimens are often part of cohorts.

## 4 Goal and Task Analysis

To understand the needs of domain experts in the field, we surveyed a group of 6 biologists and 3 pathologists from Harvard Medical School, Dana-Farber Cancer Institute, or Brigham and Women’s Hospital. From the questionnaire (see Supplemental Material) and monthly meetings over a period of one year, we identified and refined a set of high-level domain **goals** from which we derived specific **tasks** as guidelines for an effective visual analytics system. We thus fulfilled the translator role put forth in the design study methodology by Sedlmair et al. [80], requiring a comfort level with task abstraction [51] in computer science.

### 4.1 Goals

#### G1

Experts are interested in identifying how specific cell types attract or repel each other (cell-cell interaction). When immune and cancer cells are frequently observed close to one another (in each other’s spatial neighborhood), they are likely to interact. For instance, the interaction of two types of immune cells (B and T-cells) is a central tenet of protective immunity. These interactions can occur at various scales (cells directly adjacent to each other or in the same large region of tissue).

#### G2

With these neighborhoods as building blocks, experts seek to understand their spatial arrangement within the tissue image. These neighborhood patterns can be equally distributed throughout the image or appear in proximity, forming biologically meaningful spatial structures (groups). Germinal centers, which are regions within lymph nodes where B cells proliferate, are examples of such micro-structures.

#### G3

Experts seek to validate the statistical significance of identified neighborhood patterns and larger spatial structures **(G1, G2)** within and between specimens. By understanding how often they appear and how properties (e.g. composition, size) vary between patterns, expers can determine clinical relevance and motivate further investigation.

#### G4

Finally, experts want to connect patterns present in an image back to biological and clinical information. They hope to determine how the presence of specific patterns correlates to specific cancer therapies, the growth of tumors, and immune response to those tumors, with an overall goal of improving cancer diagnosis and treatment.

Our survey showed that the scientists are interested in performing both exploratory and confirmatory analysis to achieve these goals. In their highly experimental settings, discovering novel spatial neighborhood patterns and thereby generating new hypotheses is of similar importance as the ability to express and verify existing hypotheses.

### 4.2 Tasks

#### T1: Visually explore spatial neighborhoods (G1, G2)

from different perspectives. This includes navigation, visual identification, and selection of regions of interest in the tissue as a means for exploring the spatial neighborhoods present in a specimen or multiple specimens.

#### T2: Group similar cell interactions

through which experts can identify the larger structures formed by these cell-cell interaction patterns (**G2**). These grouping strategies must scale to the large data, even grouping the patterns present in multiple specimens at once while also being interactively configurable to incorporate users’ domain knowledge.

#### T3: Express and search for hypotheses (G3)

This includes the ability to query for specific user-defined cell-cell interactions as well as search by example, i.e., find additional occurrences of the pattern present in a ROI. It also includes querying across cohorts.

#### T4: Compare the contents and spatial expression of patterns

identified within a single specimen and across a cohort (**G3**), while taking into account the biological and clinical context. (**G4**). In this context, spatial expression refers to the presence of a pattern within a tissue.

#### T5: Rank the presence and statistical significance of pattern

within a specimen, building on the previous task. This augments comparison and helps experts better understand these patterns (**G1, G2**) and how they differ within and across specimens (**G3**).

#### T6: Extract, annotate, and save

found patterns along with biological and clinical information (**G4**), as analysis is ongoing and not limited to isolated sessions. This information may relate to the source of the specimen, specific treatments involved, or patient outcomes. This allows for continued analysis, where gained knowledge is applied to new specimens (**G3**), and insights can be shared with other experts.

## 5 Workflow

From the identified goals and tasks and in bi-weekly sessions with the experts, we extracted an iterative human-in-the-loop workflow (Fig. 3) for spatial neighborhood analysis that guided the design of our visual analytics system Visinity (see Sec. 7).

**Fig. 3:**
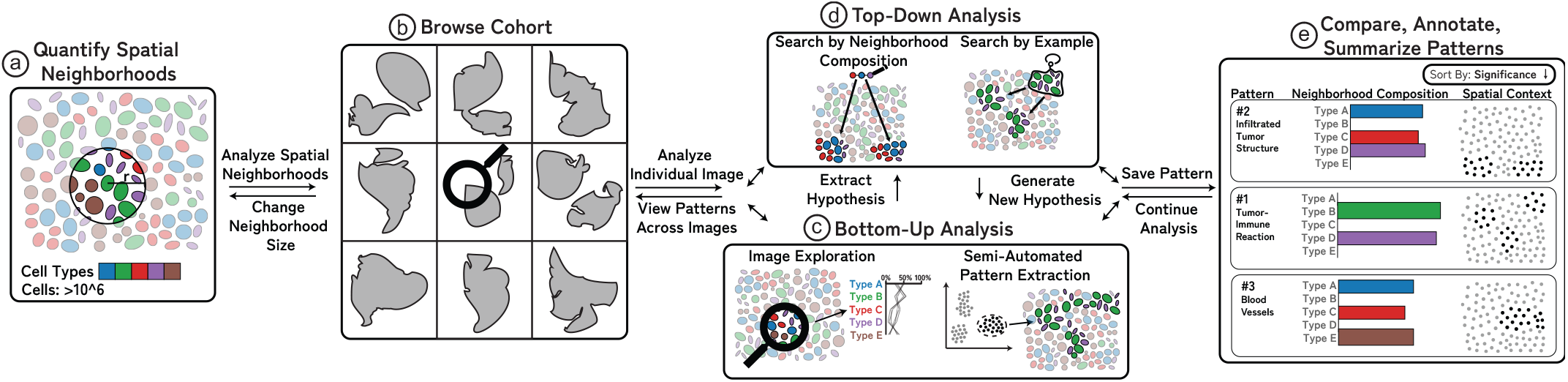
Visinity Workflow: (a) Neighborhood quantification: users choose a spatial range triggering neighborhood vector computation; (b) Browse cohort: small multiples of specimen to gain an overview of neighborhood patterns; (c) Bottom-up analysis: explore spatial arrangements and cell-type composition of neighborhoods, generate hypotheses, cluster, and extract patterns; (d) Top-down analysis: two visual querying capabilities allow hypothesis generation and search for similar patterns; (e) Pattern annotation and comparison within and across datasets.

After importing the data, users can specify a neighborhood size (can be modified), depending on if they are interested in more local or global interaction patterns (Fig. 3, a). Neighborhoods are then quantified. Users can gain an overview of the tissue environments across all specimens in their cohort **(T1)** in the form of small multiples (Fig. 3, b). Users can start to explore the image (bottom-up analysis) (Fig. 3, c) and select regions of interest in the tissue to visualize the spatial neighborhoods present (**T1**). To aid exploration, they can interactively cluster neighborhoods in an entire tissue **(T2**), extracting patterns in a semi-automated manner. Top-down analysis (Fig. 3, d, **T3**) to test and refine existing hypotheses can either be done by specifying the cell types involved in a neighborhood pattern or by selecting a region of interest in the tissue as an example of a spatial pattern. Both trigger a search within and across tissue images for matching neighborhoods. Based on the contents, spatial context within the tissue, and statistical significance of results (quantified through permutation testing), users refine their hypotheses and can save and annotate identified patterns **(T6)**. Finally, a user can (Fig. 3, e) compare these saved patterns within a dataset and across datasets, allowing them to test if a pattern in one tissue is present and statistically significant in another **(T4, T5)**.

## 6 Quantifying Spatial Neighborhoods

According to our biomedical experts, a cell’s neighborhood is defined by the cells in spatial proximity within a defined spatial distance. Groups of cells with similar neighborhoods form patterns that assemble to compose more extensive spatial arrangements. To explore **(T1**), group **(T2**), search for **(T3**), compare **(T4**), rank **(T5**), and save **(T6**) cell neighborhoods and the patterns they form, we propose a neighborhood quantification pipeline, building on existing work in the field [84, 85]. We extend these methods with a more scalable implementation to interactively investigate patterns of different length-scales in whole-slide imaging data, with spatial distance weighting to reflect the neighborhood influence of cells, and with higher-order permutation testing to determine the significance of found proximity patterns.

Our computational pipeline (Fig. 4) works in 5 steps (1-5):

**Fig. 4:**
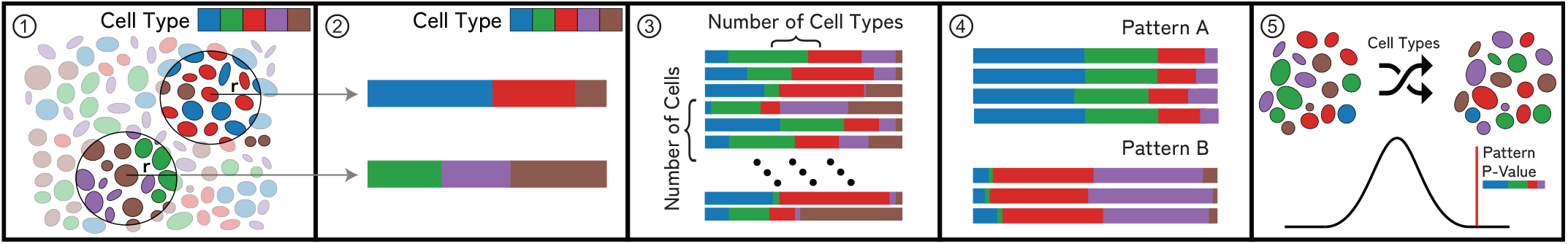
Neighborhood Quantification: (1) For a cell in an example microenvironment, find all proximate cells within a specified radius. (2) Each cell’s neighborhood is a feature vector that represents the weighted presence of each cell type in the neighborhood. (3) Repeat this process for each cell, resulting in a neighborhood vector for each cell in an image. (4) Groups of similar neighborhood vectors correspond to spatial patterns. (5) Randomly permute cell types in an image to determine patterns’ significance.

**Step 1:** We build a ball tree with the coordinates of each cell in the image. This takes *O*(*n* + *k*) to perform spatial range queries, where *n* is the number of points and *k* the number of points returned in that range [36].

**Step 2:** We create a feature vector representing each cell’s neighborhood of size 1 x *c* (where *c* is the number of cell types in the dataset). Each column corresponds to the fraction of a cell’s overall neighborhood occupied by a specific cell type. These values are linearly weighted such that cells closer to the center of the neighborhood radius contribute more to the overall neighborhood. The resulting vector is normalized. This representation builds on existing approaches [84, 85] and was reaffirmed by feedback from our collaborators, who said that it was highly interpretable and fit with the hypotheses they had regarding the cell-cell interactions present in a dataset (**T3**).

**Step 3:** Repeat this process for every cell in a dataset, resulting in one vector for each cell. We generate a matrix representing the neighborhoods in a dataset, where each row is the neighborhood of a cell.

**Step 4:** Cells with similar spatial neighborhoods are represented by similar neighborhood vectors. We run a distance-based nearest neighbor search **(T3)** and use a configurable threshold to define similarity. To find groups of similar neighborhoods **(T2)**, we utilize partition-based clustering (see Sec. 8.1 for more details). Together with our experts and based on literature [31], we evaluated vector comparison based on Euclidean distance to achieve the most satisfying results while also providing simplicity, interpretability, and scalability.

**Step 5:** Inspired by similar approaches [77, 84], we use permutation testing [30] to determine the patterns’ statistical significance **(T5)** within a specimen and across a cohort. We count individual neighborhoods that match a given neighborhood pattern based on a user-defined similarity threshold (Sec. 8.2). We then randomly shuffle the assigned cell types in the data, recomputing neighborhood vectors (Steps **1** - **3**), and calculating the number of matching neighborhoods for each permutation. Eq. 1 describes our *P* value calculation.

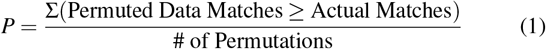

## 7 VISUALIZING SPATIAL NEIGHBORHOODS

To realize the identified workflow (Sec. 5) and task **(T1-T6)** we developed Visinity — an open-source visual analytics system [4] for spatial neighborhood analysis. Visinity’s interface (Fig. 1) consists of coordinated views offering different perspectives on detected neighborhoods, including their composition and spatial occurrence in the tissue.

### Cohort Overview

After data import and neighborhood quantification (Sec. 6), users can start the analysis with an overview of all specimens in a cohort. Together with the biomedical experts, we chose small multiples of image thumbnails as a sufficient and compact way to comparatively summarize **(T4)** the images in a cohort and their morphology (Fig. 1, b). Users can zoom and pan into these thumbnails to begin exploration and select a specific dataset for thorough investigation **(T1)** in the image viewer (a). As analysis progresses, spatial neighborhood patterns identified in a single specimen are detected and visualized in other members of the cohort by highlighting their spatial presence in linked images. Specimens can also be sorted by the number of matching neighborhoods to the pattern currently being investigated or by the statistical significance of that pattern within a specimen (see Sec. 9). **Spatial Exploration of Tissue Morphology**. To support visual exploration of the spatial neighborhoods in tissue images (**T1**), we offer a scalable image viewer (Fig. 1, a), allowing navigation via zooming+panning. The viewer builds on our previous work Facetto [40] and Scope2Screen [32]. Through multi-channel rendering, pseudo-colored channels can be blended together into a single view, enabling users to analyze the expression level of multiple markers at once.

To mark and filter for cell neighborhoods in an ROI, users can employ an interactive lasso tool. Fig. 5, a shows a selected spatial neighborhood in a lymphoid nodule from a healthy tonsil tissue. We decided to primarily render cell neighborhoods directly in the image space. This design was driven by our experts’ feedback that spatial image context is essential for pathologists to draw the right conclusions. We render selected cells with superimposed outlines so that the underlying tissue image and morphology is still visible. These outlines are optionally colored by cell type using a categorical color scale (Fig. 5, a). We derived this color scheme from ColorBrewer [28] and Colorgorical [24] to increase the contrast between selected cells and the black image background. In this example, the region is composed primarily of two immune cell types, B cells, and T cells. The encoding is effective for a few image channels and cell types (Fig. 6, a), but becomes increasingly difficult to comprehend when combining multiple pseudo-colored channels with categorical cell coloring. Thus, users can switch to an alternative mode that visualizes detected neighborhood patterns with a concave hull, emphasizing their unity while maintaining a non-occlusive view of the underlying image channels (Fig. 6, b).

**Fig. 5:**
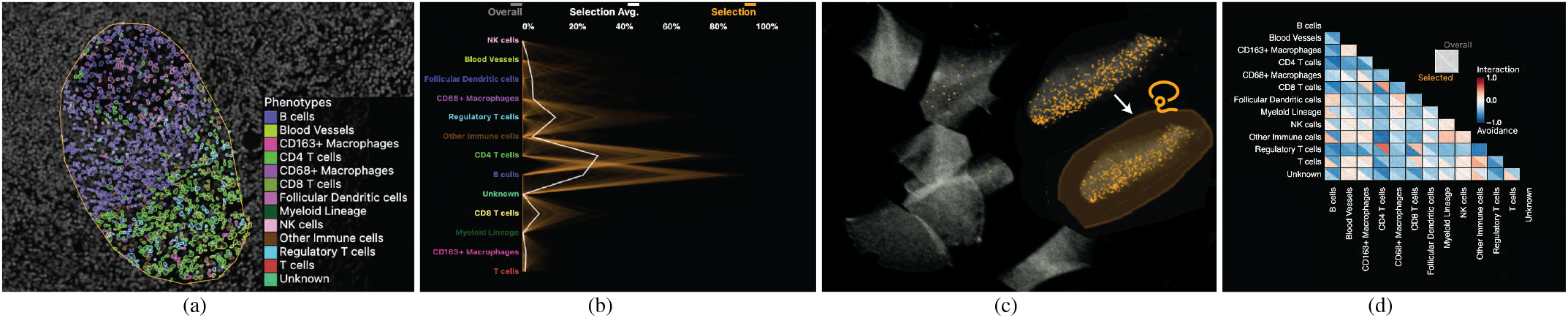
Visual Exploration Through Linked Views: (a) Selected ROI to investigate the spatial neighborhoods. Cell types are displayed with color-coded segmentation outlines. (b) Neighborhood composition in a PC plot - orange lines represent neighborhoods selected, exhibiting two discrete patterns. (c) Interactive 2D UMAP embedding of all neighborhood vectors in a specimen in grey; current selection visualized in orange. Users can select a region to explore similar neighborhoods. (d) Pairwise cell-cell interactions visualized as a correlation matrix.

**Fig. 6:**
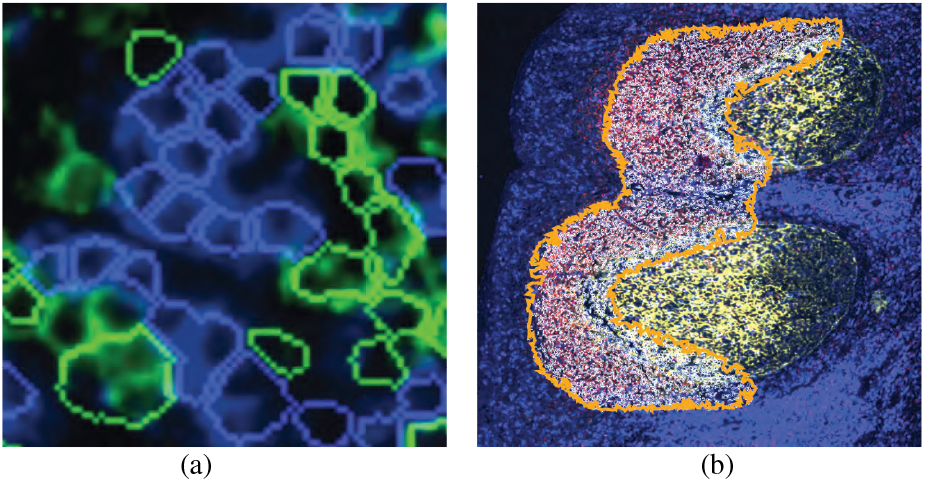
Cell Outlines or Concave Hull: Two view modes: (a) coloring cell outlines by cell type; (b) outlining patterns with a concave hull.

### Composition of Spatial Neighborhoods

To provide more information about what cell types spatial neighborhoods are composed of **(T1)**, we include a parallel coordinate plot (Fig. 5, b). We chose PC plots over bar charts and box plots to emphasize the occurrence of each cell type in a spatial neighborhood while also encoding the distribution and correlation between the features. Fig. 5, b shows the composition of a selected tonsil region (a). Here, each poly-line represents the neighborhood of a cell. Each individual axis is defined by the influence of a specific cell type in the neighborhood. Two distinct neighborhoods are represented, one containing more B cells and one containing more CD4 T cells. To emphasize correlations between cell types on adjacent axes, we employ an axis reordering strategy [9, 33]. Fig. 8, a shows a negative correlation between CD4 T cells and B cells. The axes can also be reordered by drag&drop, addressing the need of experts to investigate pairwise interactions between cell types **(T3)** To compare **(T4)** the current selection to the overall composition of a specimen or cohort, the PC plot optionally encodes the neighborhoods in the entire specimen or cohort in gray, behind the current selection in orange (Fig. 1, c). The opacity of lines in the plot is chosen based on the data size and screen dimensions, thus minimizing over-plotting and making sure neighborhood patterns within the overall data are visible. This view also supports interactive brushing, allowing users to investigate such patterns. When analyzing a specimen that is part of a larger cohort, the user can toggle between visualizing the specimen in isolation or the overall cohort.

**Fig. 7:**
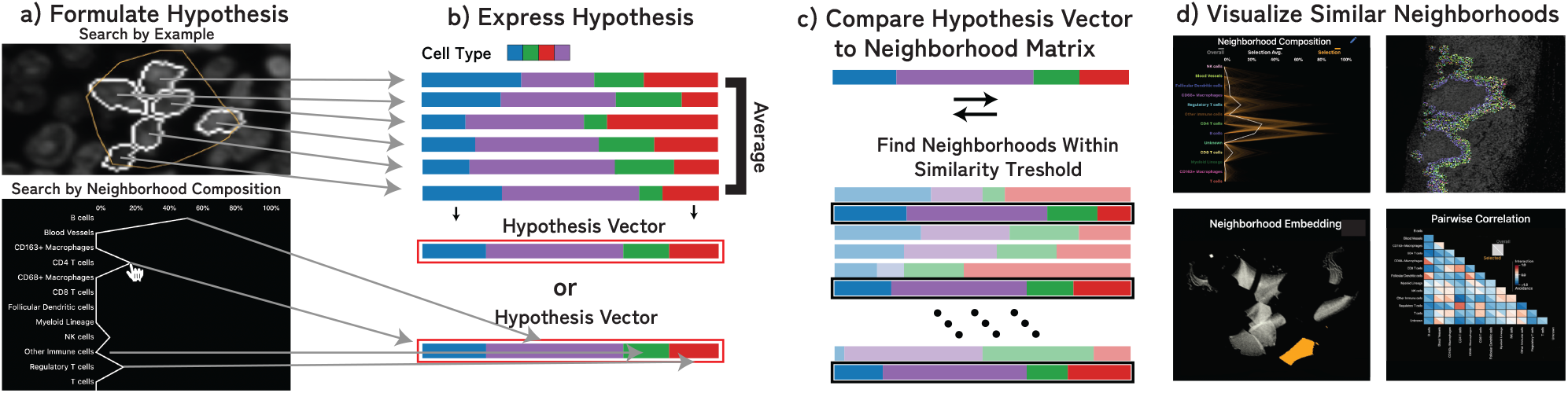
Hypothesis Testing through Visual Querying: (a) Users can test spatial pattern hypotheses in the form of regions of interest or approximate neighborhood composition. (b) We formulate this hypothesis as a neighborhood vector and (c) compare that neighborhood to every neighborhood in the image. (d) Results above a set similarity threshold are visualized in each of the linked spaces.

**Fig. 8:**
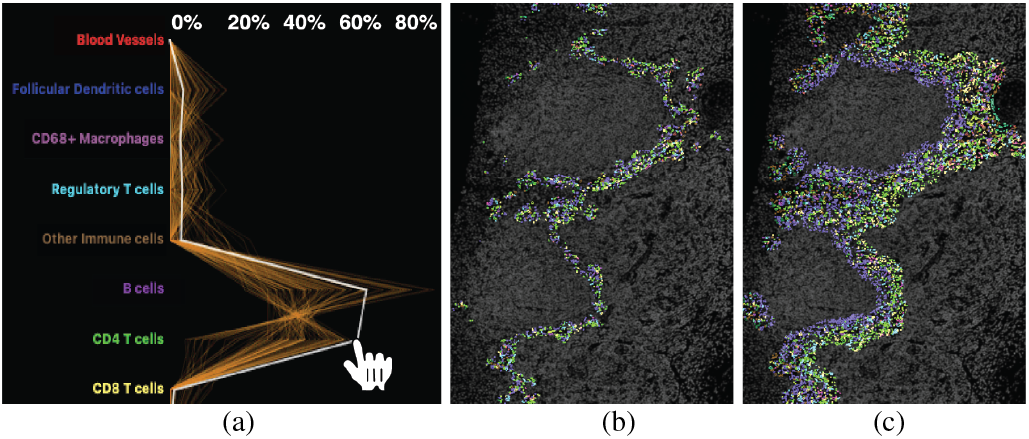
Search By Neighborhood Composition: (a) A user can sketch a custom neighborhood and increase/decrease the threshold to find more/less similar neighborhood patterns (b, c). Here, interactions between B cells (left) and T cells (right), outlining B cell follicles.

### Embedding of Neighborhood Vectors

While a PC plot emphasizes neighborhood composition and can reveal correlation, it can be hard to distinguish patterns from one another due to increasing occlusion and visual clutter. To make neighborhood patterns more distinct, we perform dimensional reduction on the neighborhood vectors and visualize the reduced 2D data in a scatterplot (Fig. 5, c), where each point reflects a cell’s spatial neighborhood. Cells that are close to another share a similar neighborhood and form spatial groups. For cohort data, we create a shared embedding of the neighborhoods across the individual datasets allowing users to identify and compare similar spatial neighborhoods and discrepancies across specimens.

Dimensional reduction is a conventional and familiar practice for our intended users. There is significant biological precedent for the use of t-SNE [38, 84, 85, 87] and UMAP [54, 85] to investigate spatial features in tissue images. With proper initialization and hyperparameters, both methods can preserve global structures and produce similar embeddings [39, 92]. While both are stochastic, UMAP has been shown to demonstrate improved stability, making the embedding more reproducible [54]. We thus use UMAP with parameters to emphasize global structure [15] (50 n neighbors, 0.01 min dist) to visualize these data, though t-SNE with a high attraction behaves similarly [11]. We found that UMAP produced good spatial separation in the 2D layout at various dimensionalities (dictated by number of cell types), allowing users to distinguish patterns from another and easier selection. The scalability of UMAP is another benefit, specifically when using RAPIDS GPU UMAP implementation [59], which is capable of embedding million-cell neighborhood matrices in a few minutes (see Sec. 10). However, we see incorporating other dimensionality reduction techniques, particularly those which emphasize scalability [37, 66], allow for user input [65], or are tailored to specific biological data (e.g. preservation of rare cell types [88]) as a promising application for Visinity (Sec. 11.4).

Users can navigate in the scatterplot (embedding view) and make selections as they would with the image view, which highlights the selection in the coordinated views. E.g., a user may notice a region of high density in the embedding (Fig. 5, c) and select it in order to understand the cell types that compose that visual group (Fig. 5, b) and the locations (Fig. 5, a) of those neighborhoods within a tissue image (image viewer) or across multiple images (cohort view). Likewise, users can select a spatial region in the cohort view and image viewer and review the greater neighborhood pattern it is part of in the embedding. When investigating a specimen, users can toggle between the individual embedding or cohort embedding.

### Pairwise Correlation Between Cell Types

To address the need to explore pairwise interaction between two specific cell types **(G1), (T1)** we offer a correlation matrix visualization. We chose a matrix over a node-link diagram to avoid clutter when encoding relationships between every pair of cell types. Matrix visualizations provide a consistent and compact layout, making it easy for users to compare interactions in a specific pattern to those in the overall specimen and to other patterns **(T4)**. Inspired by existing geographic [52] and biomedical [85] approaches, we compute the Pearson correlation coefficient between each pair of cell types within the computed neighborhood vectors to quantify these pairwise relationships. Two cell types with a strong positive correlation tend to be found in similar neighborhoods, whereas two cell types with a strong negative correlation tend to avoid each other. We use a diverging red-white-blue color palette to visualize these correlation values. To support comparison, we split each field into two triangles (Fig. 1, e), one representing the correlation of cell types in the overall image or cohort and one representing correlations within currently selected neighborhoods. Selecting a triangle in the matrix filters the other views to highlight neighborhoods with respective pairwise cell type correlation above a user-defined threshold. In Fig. 5, d, B cells have a negative correlation to all other cell types (all correlations are blue), indicating that the region of B cells is very homogeneous. CD4 T cells, meanwhile, have a positive (red) correlation with CD8 T cells, blood vessels, and regulatory T cells, indicating these different cell types are interacting.

## 8 Semi-Automated Analysis

Beyond exploration through linked views **(T1**), Visinity offers semi-automated methods to cope with the large and high-dimensional (*≥*million cells per specimen) data. This allows users to automatically group spatial neighborhoods **(T2)** through interactive clustering and search for neighborhood patterns to test and refine hypotheses **(T3**) at scale.

### 8.1 Detecting Spatial Patterns Through Clustering

To automatically cluster the cells based on similar neighborhood vectors we chose to use EM (expectation-maximization) clustering for Gaussian mixture models [96]. Depending on the specific biomarkers, derived cell types, and cellular neighborhoods, we can only make assumptions about the underlying model with many latent unknown variables. The EM algorithm finds (local) maximum likelihood parameters of that statistical model given our sample data [16] and can detect clusters that vary in shape and density, compared to, e.g., k-means. This approach scales to cluster our datasets interactively without precomputation.

The interface enables users to either cluster the neighborhoods in a specific dataset and apply this clustering to the rest of the cohort, or to run the clustering on all specimens in a cohort and then drill down into individual specimens to explore results in more detail. Computed clusters are listed with other saved patterns in a list view (see Fig. 1, i), and users can click on them to visualize them in each of the linked views. The clustering can be used in an iterative manner; based on the spatial context in the image, the cluster’s composition, and where the cluster lies in the embedding, a user may choose the number of clusters they desire, allowing the detection of sub-structures that may exist within a given neighborhood pattern as well as the macro-structures that contain these neighborhoods compose. We visualize these clusters by coloring their cells by cell type and emphasize unity with a concave hull, as we do for regional selections. Fig. 10 shows identified clusters in the tonsil data, including the B cell follicles investigated earlier and the *‘Paracortex’*, containing many different types of T cells.

**Fig. 9:**
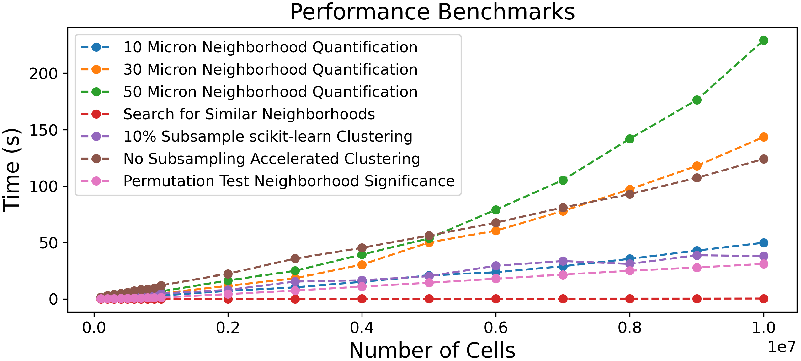
Runtime evaluation for steps in the neighborhood computation pipeline. Data size is increased gradually.

**Fig. 10:**
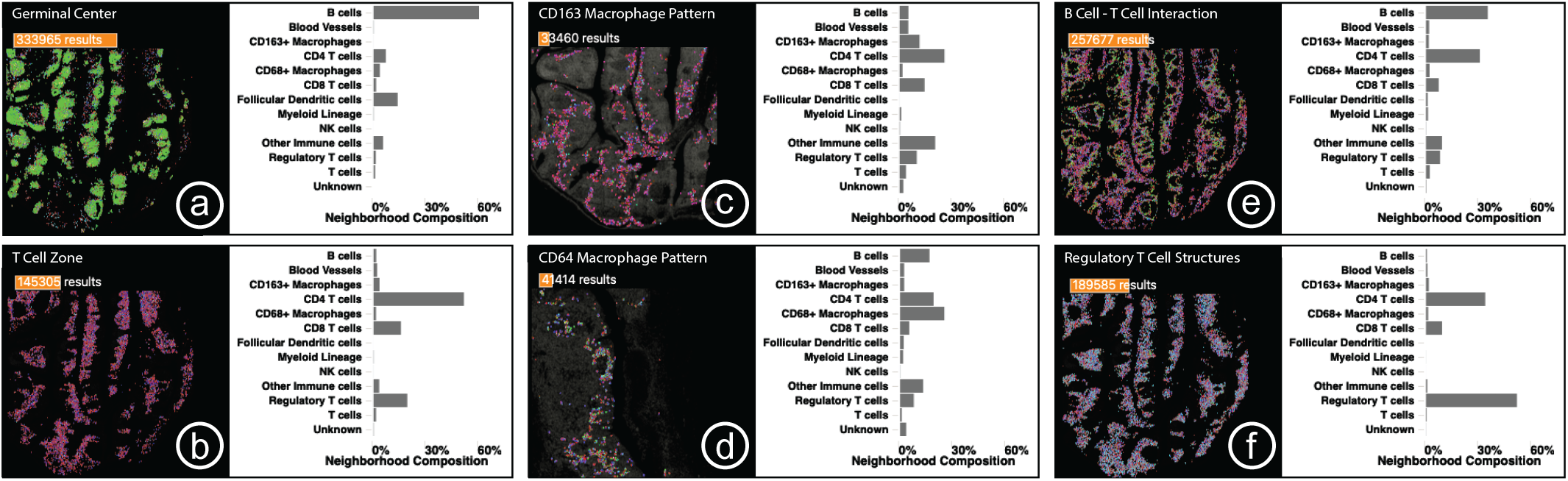
Case Study 1: Small multiple comparison views summarize the biological structures and processes found in a human tonsil, including key interactions between B cells, T cells, and macrophages as well as unexpected regulatory T cell structures.

### 8.2 Hypotheses Testing With Visual Querying

#### Visual Querying By Region of Interest

For any region of interest found during data exploration, there might be similar regions in the specimen or throughout the cohort. Visinity enables users to execute such queries for any detected or manually selected neighborhood pattern (query-by-example), revealing similar spatial neighborhoods both within the image and in all images in the cohort (Fig. 1, g).

The computational process is outlined in Fig. 7: For any selected region (a), we first compute the mean of the neighborhood vectors in that region (b). We then compare this representative vector *v*_1_ with each other neighborhood vector *v*_2_ in the image using the Euclidean similarity score [81]: 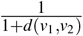, with distance *d* between vectors. Neighborhoods above a specified similarity threshold are then highlighted in each of the linked views (Fig. 7, d). Additionally, we visualize (Sec. 9) the number of matching neighborhoods and statistical significance of this hypothesis as determined through permutation testing (Sec. 6).

Representing an ROI by its mean vector is an approximation of the neighborhoods in the region. As the region increases in size, particularly when the region contains multiple patterns of interest, this gets more and more inaccurate. However, when such a region is selected, the composition and embedding views provide context to the homogeneity of this selection, allowing a user to investigate potentially discrete patterns in isolation. Moreover, to accommodate the analysis of regions of tissue of any size, users can modify the neighborhood radius to fit their needs. Through iterative design with frequent feedback from our expert collaborators, we found this approach was effective at finding similar neighborhoods, as demonstrated in Sec. 11.1 and Sec. 11.2.

#### Visual Querying By Neighborhood Composition

Experts indicated that they often were interested in investigating interactions between specific cell types **(T3)**. To support expression and search for their hypotheses (Fig. 5, f), users can look for specific interactions by defining a custom query vector in the neighborhood composition view by clicking and dragging to create a polyline representing a query vector. The current neighborhood composition and overall composition are both optionally displayed behind the query vector, allowing the user to match trends reflected in these views while building a query (Fig. 8, a). The results, which represent the interaction of the specified cells, are visualized in the linked views (Fig. 8). A user can perform query refinement or expansion by adjusting the similarity threshold; increasing this threshold yields neighborhoods that more precisely match the query (Fig. 8, b) while decreasing the threshold casts a wider net (Fig. 8, c). The same refinement can be applied to ROI-based queries.

### 8.3 Visual Querying Across Specimen

When a user queries a specimen for a pattern, we search across the entire cohort for matching neighborhoods, which are displayed in the cohort section of the comparison view (Fig. 1, f). We additionally display the number of results and significance of the query within each specimen above the spatial summary, as discussed in Sec. 9. Users sort related specimens within the list by significance and number of results **(T5)**.

## 9 Visual Comparison and Result Summarization

While using Visinity, users can save and retrieve patterns and label them with clinical or biological context **(T6)**, which are displayed in a pattern list in the interface, as reflected in the proposed workflow (Fig. 3, e). Once saved, experts can compare the neighborhood composition and spatial expression of these patterns **(T4)**. Comparison can happen on two levels: between identified patterns and between specimens.

### Comparison Between Neighborhood Patterns

To summarize and compare identified neighborhood patterns, users can choose between spatial and compositional comparison (Fig. 1, f). Small multiples of bar charts (Fig. 10) represent the composition of each saved pattern, allowing users to compare the presence of each cell type between patterns. We chose bar charts over the initial PC plot (Sec. 7) as they are a concise and simple visual encoding that allows users to easily compare the neighborhood composition of many patterns **(T4)**. We also provide spatial context regarding the patterns in a concise manner with scatterplots (Fig. 10), which encode the position of each pattern in the tissue, providing a stripped-down visual summary of the tissue imaging data. A user can select these small multiples to view them in full detail in each of the linked views and, in the case of comparison across images, to search for that pattern within a new image. On top of each plot, a bar represents the number of cells with that given neighborhood above each specimen thumbnail (Fig. 10). The color of the bar encodes the statistical significance (see Sec. 6, step 5) on a single hue white-orange color scale. **Comparison Between Specimens**. It is also of interest to understand where and how frequently a specific neighborhood pattern appears in the different specimens **(T4)**. After a pattern is selected from the pattern list (Fig. 1, i), a bar shows the number of matching neighborhoods above each specimen thumbnail of the Cohort View (Fig. 10). Again, we encode the number of cells contained in the pattern with its computed significance as a bar above the respective plot.

## 10 Scalable Implementation

Throughout the design process, we emphasized scalability, leveraging methods and interfaces that support simultaneous analysis of gigapixel images containing more than a million cells. We feature a JavaScript client / Python server architecture with web-based frontend visualization and efficient backend computation. We built on our previous work (Scope2Screen [32], Minerva [29, 70]), storing images in the Zarr [55] format and rendering them using WebGL. The embedding view, linked image thumbnails, and spatial comparison views use the regl-scatterplot [43] library, which allows for panning, zooming, and selection in datasets containing as many as 20 million points. We combined efficient methods with strategic pre-computation to quantify and analyze spatial neighborhoods. Neighboring cells are queried using scikit-learn’s [63] ball-tree index structure. The neighborhood vector computation is compiled with Numba [41], and thus translated into efficient machine-code. When the neighborhood radius changes, we recompute all neighborhoods and save them as Zarr arrays. To evaluate runtime performance of this approach (Fig. 9) we generated random test data ranging in size from 10^5^ cells to 10^7^ cells. These synthetic data maintain the same cell density as the Tonsil dataset investigated in Case Study 1. We randomly give each cell one of the 13 cell types assigned in the same tonsil dataset, maintaining the same incidence of each cell type as was present in the tonsil. The greatest computational bottleneck is neighborhood quantification at increasingly large neighborhood sizes, which is primarily a result of the time needed to identify neighboring cells. We found the ball tree used to search neighbors outperformed various other similarity search implementations [26, 34] on 2D data. When computing neighborhoods, we also randomly reassign cell types to create new permuted versions of this neighborhood matrix, which is similarly compiled and saved as a compressed Zarr array. Similarity search across the neighborhood vectors and permuted neighborhood vectors is conducted in parallel. We offer two EM clustering implementations, one using scikit-learn [63] which is fit on a 10% random sub-sample of the data and a hardware-accelerated approach [10] fit with all the data. Sub-sampling worked well with data and quantified cell types used by our collaborators. However, when small quantities of rare cell types are identified, these cells may be lost. SCHNEL [5], among other methods offer clustering strategies that specifically preserve rare cell types.

As demonstrated in Fig. 9, our permutation testing, clustering, and search implementations scale linearly as the number of cells increases. Visinity’s source code and executables are available on GitHub [4].

## 11 Case Study Evaluation

We present two case studies that demonstrate the utility of our system, each with a domain expert. These experts were involved in the goal and task analysis and provided incremental feedback during the system development. In each 90-minute in-person session, the participants, who had no hands-on experience with Visinity, were given a brief walk-through and then steered the system themselves. While analyzing the tissues, the experts were instructed to think-aloud the biological context and motivation for their analysis and to provide usability feedback for the system. After completing the session, each expert filled out a survey quantifying the usefulness and intuitiveness of Visinity’s features.

### 11.1 Case Study 1: Human Tonsil

Tonsils, which are lymphoid organs at the back of the throat, are a part of the immune system and help defend against foreign organisms. They have been extensively studied as they are dense tissues with distinct morphology and contain many immune structures and interactions. In this case study, a senior anatomic pathologist (P1) at Harvard and Brigham and Woman’s Hospital, with a focus on precision medicine in cancer biology, investigated a single tissue specimen of a normal human tonsil. We demonstrate that Visinity can be used to identify such known spatial arrangements of immune cells.

#### Data

The tonsil was scanned to generate a 17,231 × 12,312 micron whole-slide image of 1.3 million cells. The imaging data are 62 GB and contain 45 channels, each 26,482 × 19,065 pixels. Cells in the specimen were assigned by a computational biologist to one of 13 different types based on patterns of protein expression.

#### Analysis

We began the analysis by turning on the DNA channel in the image viewer, outlining the nuclei of all cells in the specimen in the tissue image. Zooming and panning in the image revealed the expected tonsil morphology and architecture. As an initial exploratory step, we clustered the spatial neighborhoods in the tonsil into ten groups. P1 immediately noticed three clusters occupying adjacent regions in the image that corresponded to known biological structures and interactions. One cluster formed distinct oval-shaped groups, which P1 identified as representing germinal centers which are areas of B cells, a type of white blood cell that proliferates before migrating out of the tonsil to secrete antibodies (Fig. 10, a). Another cluster outside of the germinal centers contained numerous T cells of various types (helper, cytotoxic, and regulatory) and specialized blood vessels that shuttle B and T cells in and out of the tonsil. P1 identified this as the T-cell zone of the tonsil (Fig. 10, b). Between these clusters was a band of Helper T cells binding to B cells. Here, Helper T cells facilitate in the development of B cells into antibody-secreting plasma cells or memory B cells, which remember information about specific antigens so the body can defend against them in the future (Fig. 10, e).

P1 then investigated the distribution of macrophages in the tonsil, which are cells that engulf and consume other cells and attract immune cells. In this tissue, two types of macrophages were identified based on the proteins they express, CD68 and CD163. We selected each type of macrophage in the cell-type legend and found that these neighborhoods occupied two distinct regions of high density in the embedding. We saved and labeled both patterns. In the spatial comparison view, the pathologist noted that the CD68 macrophage neighborhoods were located in the interior of the tonsil, near and within the B cell follicles, whereas CD163 macrophages were closer to the surface of the tonsil (Fig. 10, c,d). The comparison bar charts demonstrated that while both macrophage neighborhoods contained roughly the same fraction of Helper T cells and Regulatory T cells, the CD68 macrophage neighborhoods were far richer in B cells, whereas the CD163 macrophage neighborhoods had more T cells and fewer B cells, suggesting differential roles in B cell development.

In addition to identifying common immune interactions and distributions, the analysis revealed less well-appreciated patterns. The correlation matrix showed that Cytotoxic T cells and Regulatory T cells had the highest correlation in the specimen. We selected this index in the matrix to find neighborhoods of both cell types. Regulatory T cells help prevent the immune system from attacking healthy cells and the pathologist noted that the co-localization with Cytotoxic T cells is consistent with a modulatory role for the regulatory cells in controlling cytotoxic T cells, which kill other cells. In the composition view, most of the neighborhoods were shown to contain a roughly equal proportion of the two cell types as well as some helper T cells, but some neighborhoods were visible with a higher incidence of Regulatory T cells. Sketching such a pattern and searching for similar neighborhoods revealed small groups of cells underneath the tonsil membrane and outside of the B cell follicles (Fig. 10, f). P1 indicated that it was interesting to find Regulatory T cells clustered together since they are generally more evenly dispersed throughout tissues.

### 11.2 Case Study 2: Lung Cancer in Mouse Tissues

Mice share many biological similarities with humans and are commonly used as a model organism to study cancer biology. During this case study, a senior biologist (B1) with expertise in quantitative, molecular, and cellular biology at Harvard and Brigham and Women’s Hospital investigated 10 lung tissues belonging to a cohort of mice that were genetically modified to develop multiple small tumor nodules. Within this cohort, half of the specimens were from mice that develop tumors that elicit a very poor immune response (immune-poor) while the other half were engineered to activate Cytotoxic T cells, driving a robust infiltration of immune cells into the tumors (immune-rich). The objective of this study is to identify spatial organization and molecular features of immune cells that prevent tumor growth and to characterize the features of the immune response between the two specimen types.

#### Data

This cohort was composed of 10 specimens representing 740 GB of imaging data. Each image contains 31 channels and is more than 600 million pixels in size. In total, these specimens contain 2.6 million cells, each assigned to one of 17 cell types by the senior biologist.

#### Analysis

As lungs contain many regions with low cell density, we chose a wide 50-micron neighborhood radius. B1 began in the cohort view to get an overview of each specimen and its morphology and then looked at the cell types with the highest pairwise correlation across the cohort in the matrix visualization, which were Dendritic cells and T cells. These cells are known to interact; Dendritic cells are messengers, presenting antigens to the surface of T cells, activating T cells to combat tumors. B1 indicated that identifying this interaction confirmed the accuracy of the cell-typing.

B1 next investigated the neighborhood embedding and changed the plot to color by cell type; this revealed a region containing many different immune cells, which was confirmed by selecting the region and visualizing these neighborhoods in the composition view. Adjoining this region in the embedding, the biologist identified a large cluster composed of a high percentage of epithelial cells. They selected one half of the epithelial region of the embedding; when visualized spatially across all specimens, this region corresponded to epithelial cells that make up the airways in the lung. The other half was identified as containing many of the tumors. B1 clicked on a specimen in the cohort view to investigate it individually. Turning on the channel for TTF1, a biomarker used to recognize lung tumors, allowed the biologist to identify many separate tumor nodules throughout the tissue, two of them in close proximity to each other. After lassoing one of the tumors, the outlined cells and composition view showed mostly epithelial cells, as well as a small number of macrophages and Cytotoxic T cells on its periphery. B1 identified this tumor as not infiltrated by immune cells. We used neighborhood similarity search to identify this pattern across both this single specimen and the entire cohort of specimens, identifying several other structures within the specimen that the biologist (B1) determined to be immune-poor tumor. When assessed across the cohort and sorted by the number of matching neighborhoods, the biologist found that far fewer of these structures were identified in the immune-rich cohort than in the immune-poor cohort, consistent with the lack of tumor antigen expression and immune activation in the immune-poor cohort. We investigated these tumors in the image, B1 noticed many adjacent immune structures, which occupied a distinct region in the embedding space. This region corresponded (Fig. 11, a) to immune structures outside of tumors across the specimens. Returning to the initial specimen we were investigating, B1 selected a tumor in the image which did not belong to the non-infiltrated tumor pattern. This pattern occupied a distinct region in the embedding space (Fig. 11, b), which, when investigated in all specimens, we found to represent immune-infiltrated tumors. The linked views demonstrated that the selected tumor contained many B cells as well as Cytotoxic and Helper T cells. Despite arising in an animal that was not engineered to express a tumor antigen, this tumor was infiltrated by immune cells, highlighting the natural variability present in animal models of disease.

**Fig. 11:**
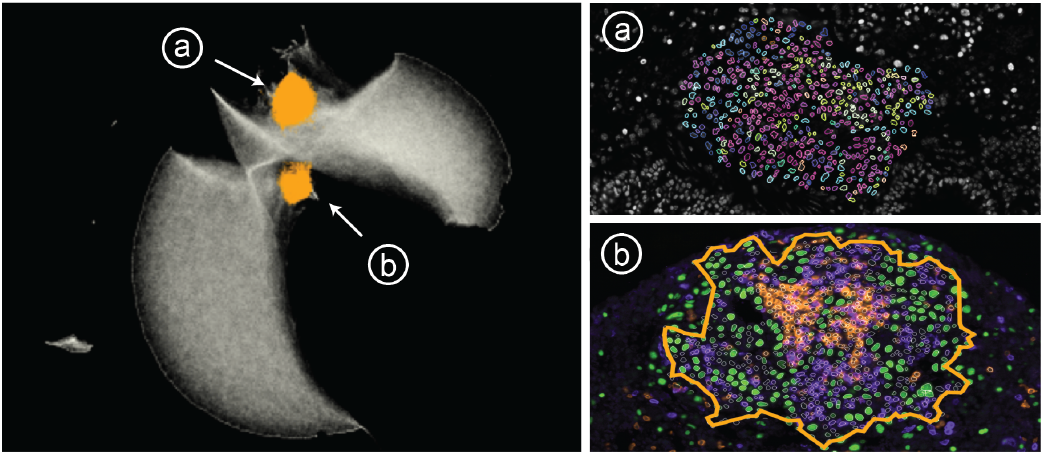
Case Study 2: Distinct regions in the embedding represent (a) immune structures far from tumors and (b) tumors infiltrated with immune cells. Tumor cells (green), B cells (orange) and T cells (purple).

### 11.3 Feedback and Survey

We collected think-aloud feedback [12] from the pathologist (P1) and biologist (B1) during the case studies. We also held additional 30-minute hands-on sessions with two biologists at Harvard Medical School not involved in the project: a postdoc and a research assistant, both with expertise in the tumor microenvironment and multiplex immunofluorescence imaging. The users then filled out a survey to rate Visinity’s features on a 5-point Likert scale ranging from strongly disagree to strongly agree.

All users rated the interface design as intuitive and accessible (4x strongly agree). They particularly liked the search by region of interest (4x strongly agree) and by composition (4x agree). Both P1 and B1 emphasized that searching for known cell-cell interactions is a powerful way to verify cell types and confirmed the effectiveness of the similarity search in various cases. While all users strongly agreed that the matrix helps to identify co-localization of cell types and can highlight pairwise interaction patterns, P1 indicated that the lack of stronger correlations and, thus light shades of blue or redmade it more difficult to interpret. The composition view was rated helpful and intuitive (3x strongly agree, 1 agree). B1 suggested allowing users to switch between displaying the relative and normalized presence of a cell type in neighborhoods on the x-axis of the PC plot; some types occurred infrequently in the mouse cohort, making it hard to compare their incidence in neighborhoods.P1 particularly liked switching between user exploration and cross-sample testing. and suggested adding functionality to facet specimen into individual sets, allowing for regional comparison in a similar way. When comparing patterns across specimens, users generally found that patterns were either wholly statistically significant or insignificant. B1 noted that while permutation testing is the standard way of determining significance, other statistical approaches that did not assume that the axis of variability was similar to the axis of information might yield more nuanced results. B1 stated that moving from a single sample to groups of samples made Visinity uniquely suited to robust and reliable biological discovery and was not included in any other visualization tool they had been exposed to.

### 11.4 Lessons Learned

Long-term collaboration with biomedical experts was essential to understand the application domain. The digital pathology field is familiar with visualization, e.g., ways to display multiplexed image data, colorization of channels, projections of high-dimensional features, and heatmaps (matrices) to discover cell interaction. Learning the experts’ dictionary and conventions was key to providing a useful solution while offering concepts beyond the state-of-the-art. Together, we refined features and designs from early and separate prototype views into integrated solutions. We identified questionnaires (to understand the needs), real-world use cases (to test the applicability), and hands-on user testing (to stress the interface design) as a successful combination for our evaluation. Secondly, we learned and categorized approaches to quantify spatial neighborhood depending on the data and application scenario, but we experienced a lack of consensus and integration. A VA approach offering round-trip analysis rendered highly advantageous for them compared to the state-of-the-art of disconnected tools for individual steps. Through iterative design, we discovered a tight linkage between image, composition, and embedding perspectives a powerful analysis concept. We found that displaying neighborhood patterns in the image with minimalistic outliness and boundary encoding to be an effective approach aligning with our experts’ expectations and existing conventions. We identified ‘cellular neighborhood’ and ‘cell interaction’ as constructs of spatially close cells and ‘spatial neighborhood patterns’ as regions of interest with frequent occurrences of these constructs (building blocks). We learned extending visual neighborhoods analysis from a single specimen to evaluating neighborhood patterns in cohorts is key to discovering significant building blocks in the cancer micro-environment.

## 12 Conclusion and Future Work

We present Visinity, a visual analytics system for investigating spatial neighborhood patterns within and across tissues. It provides a flexible human-in-the-loop workflow building on an integrated computational pipeline to quantify cellular neighborhoods. We demonstrate the applicability of our system to identify biologically meaningful spatial neighborhood patterns. We identify three key avenues of future research, ranging from the short to the long term.

### Extracting Image-Based Features

Our methods for quantifying and identifying spatial patterns are built for single-cell information (position and cell type). However, reducing high-detail images of cells to single intensity values or type classifications causes a loss of information. Computer vision models could help to capture biological structures based on their shape, as well as marker polarization within cells, indicating if cells attract or repel each other. While many deep learning models for image-based feature extraction are a black box, visual analytics could add interpretability and steerability.

### Moving Beyond 2D Imaging

Recent developments in tissue imaging have begun to produce multiplexed high-resolution 3D volumes. While our scalable method for quantifying neighborhoods adapts well to 3D data, such multi-volumetric data pose new challenges in designing suitable visual encodings, scalable volume+surface rendering strategies, constraint and guided navigation, and interaction.

### Identifying Spatial Signatures

As spatial analysis of tissue imaging data becomes more prevalent, it would be helpful to extend Visinity’s existing ability to save significant spatial patterns into a knowledge database similar to the Molecular Signature Database [47] that serves as a vital reference in genomics or even assemble the relations into a knowledge graph representation as provided in the INDRA project [27]. Accompanying visual interfaces to query and explore attributes and relationships of biological relevant spatial interaction patterns could greatly benefit the digital pathology community, allowing the discovery of new patterns and easing communication of cancer research.

## Acknowledgments

This work is supported by the Ludwig Center at Harvard Medical School, NIH/NCI grant U54-CA225088, and NSF grants IIS-1901030 and NCS-FO-2124179.

