## Supplementary Material 1 for "Visinity: Visual Spatial Neighborhood Analysis for Multiplexed Tissue Imaging Data"

Expert Survey (Questionnaire Results after Hands-On Usage of Tool)

| | Strongly disagree | Disagree | Neural | Agree | Strongly Agree |
|---|---|---|---|---|---|
| **Visual Encodings** | | | | | |
| Displaying color-coded outlines around cells indicating their cell type shows which cells may be interacting. | | | | 1 | 3 |
| Visualizing spatial neighborhood patterns with the parallel coordinates plot helps show which cell types cellular neighborhoods are made of. | | | | 3 | 1 |
| Visualizing a 2D embedding of spatial neighborhoods as a scatterplot helps uncover cells with similar spatial neighborhoods and spatial neighborhood patterns. | | | | 1 | 3 |
| The matrix visualization helps to identify colocalization of pairs of cell types and can highlight interesting interaction patterns. | | | | | 3 |
| **Discovering and Searching For Neighborhood Patterns** | | | | | |
| Searching for similar neighborhoods based on their composition (e.g. 50% Cell Type A, 25% Cell Type B, 25% Cell Type C) helps test hypotheses about the spatial neighborhood patterns within a specimen or cohort. | | | | | 4 |
| It is easy to sketch a neighborhood in the composition view (Parallel Coordinates Plot) to express a spatial neighborhood hypothesis. | | | | 1 | 3 |
| Searching for similar neighborhoods based on a selected region of interest can uncover spatial neighborhood patterns within a specimen or cohort. | | | | | 4 |
| It is easy to use the lasso tool to select a region of interest in the tissue image. | | | | | 4 |
| Using Visinity's clustering functionality helps quickly identify neighborhood patterns in a specimen or cohort. | | | | 2 | 1 |
| The interface for clustering is intuitive. | | | | 1 | 3 |
| **Saving, Labeling, and Comparing Patterns** | | | | | |
| It is helpful to keep track of found patterns by saving and labeling them. | | | | | 4 |
| The interface for saving and labeling patterns is intuitive. | | | | | 4 |
| Small visual summaries indicating where in a specimen a spatial neighborhood pattern exists help compare this pattern to others. | | | | 1 | 3 |
| It is useful to compare patterns by looking at which cell types they are composed of. | | | | 2 | 2 |
| **Overall Impressions** | | | | | |
| The application interface design is intuitive and accessible. | | | | 2 | 2 |

**Is there any other feedback you can share?**

The tool is easy to use and provides a number of features that enable both testing spatial biology swiftly switching between user exploration and cross-sample testing. The visualization interface that allows moving from single sample to groups of samples is intuitive and extremely useful - it is not included in any data visualization tool I have been exposed to and it make Visinity uniquely suited to robust and reliable biological discovery.

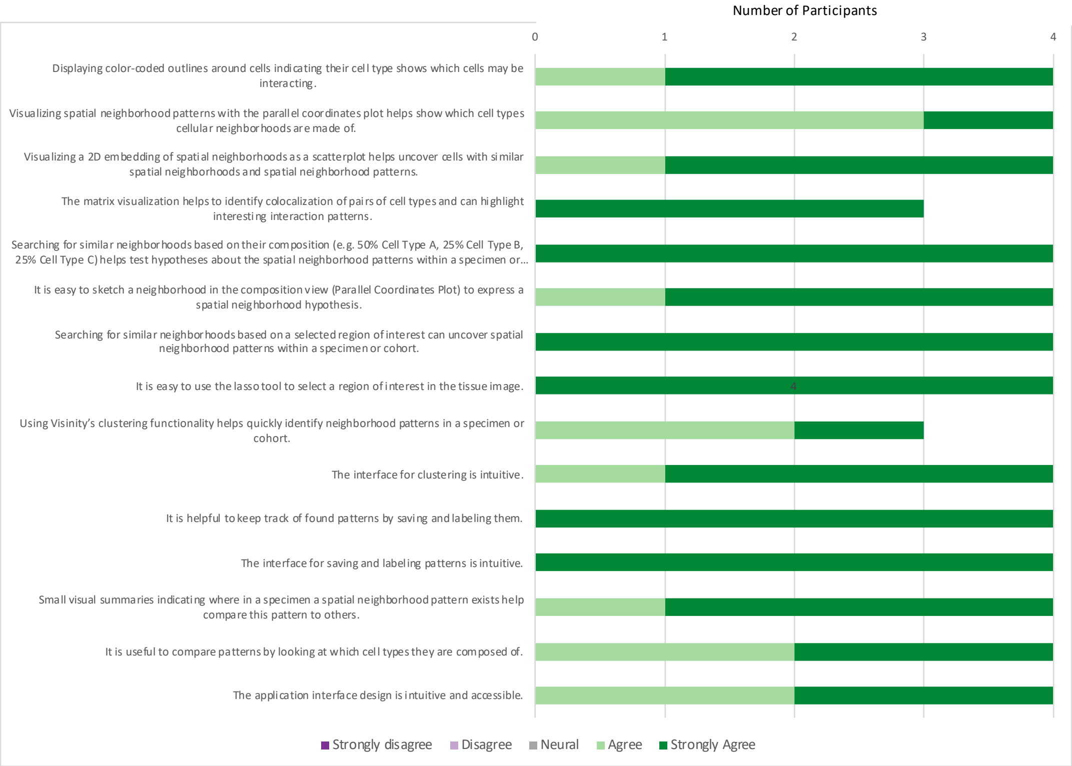
