## Supplementary Material 2 for "Visinity: Visual Spatial Neighborhood Analysis for Multiplexed Tissue Imaging Data"

### Neighborhood Analysis – Expert Feedback

6/27/2022

This document is supplemental material to the IEEE VIS 2022 submission:

We surveyed 9 experts in the field of digital histopathology. The survey was carried out online using Google forms. This document displays the answers of the participants in textual and graphical (aggregated) form. Names and other personal information contained in answers were removed. The order of the participants' answers varies by question. These measures were taken to keep the participants anonymous.

*We received an IRB (Institutional Review Board) approval from the committee on the Use of Human Subjects at Harvard University as well as the participants' acknowledgments to carry out and publish this survey.*

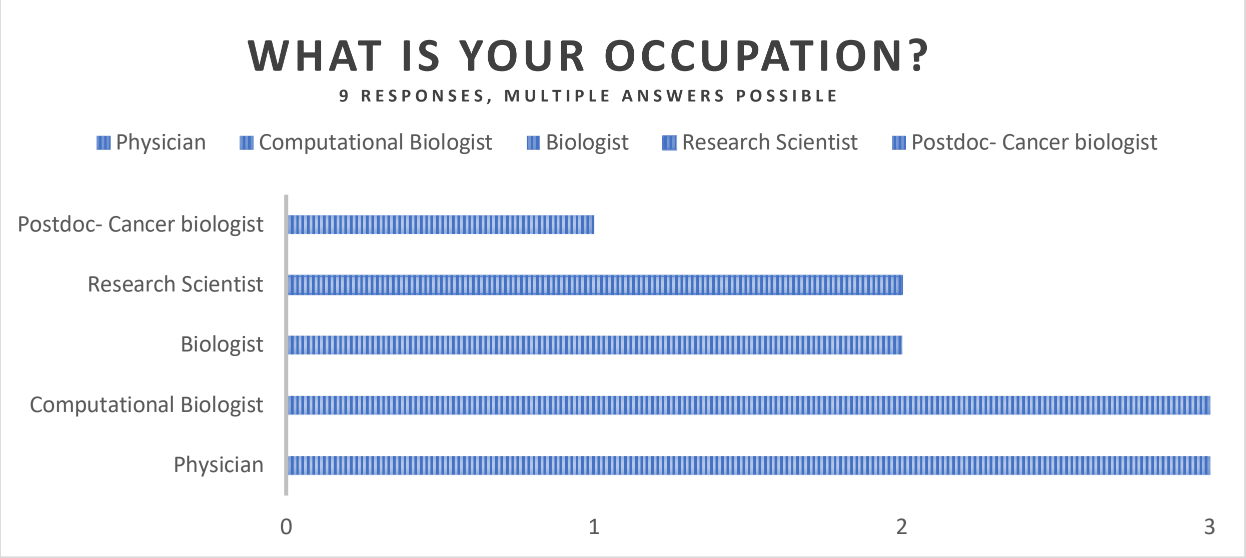

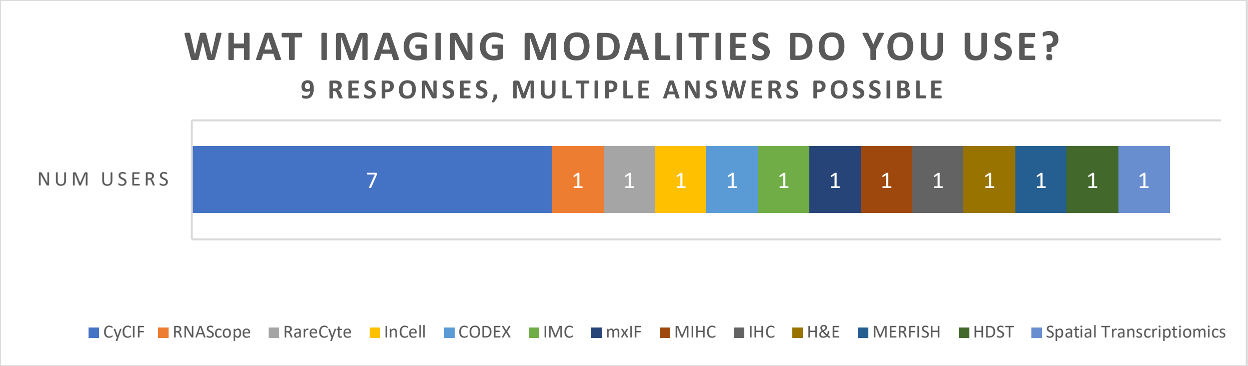


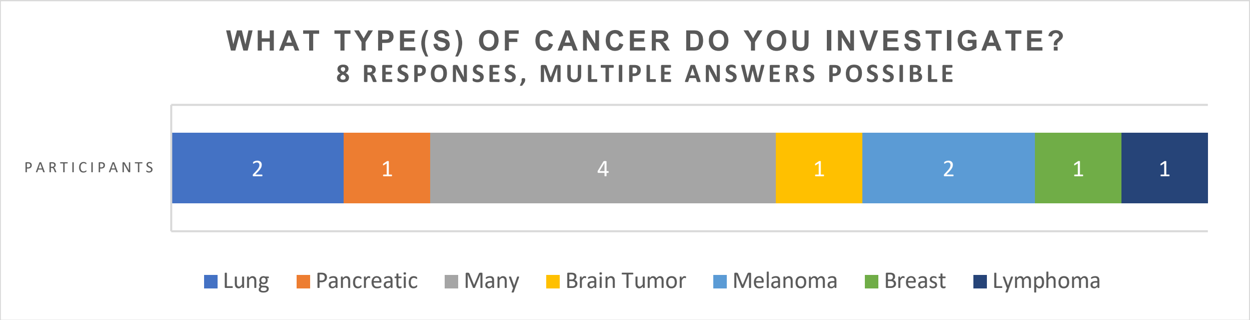


**Can you provide specifics about the size and scope of your datasets? How large are they (# of cells, pixels, storage size) and how many markers do they generally contain? (Whatever specifics you know off the top of your head are fine)** **9 responses**

| |
|---|
| 20-30 channels and around 1 million cells |
| Single tissue sections are typically 10^4 - 10^6 cells. Often 20-40 markers excluding DNA-stains. |
| Millions of cells per dataset, up to 100G per image, >100 million pixels ; >30 markers |
| >1,000,000 cells per dataset (>500M pixels per images); 20-30 markers/images per samples |
| WSI (~2-6cm2), 20-30 markers; large gigapixel images when registered and stitched; gigabyte to terabyte datasets |
| The majority of my work is focused on the development and implementation of new algorithms. Thus, most of the data I work with are exemplar and benchmark datasets. Recently, we introduced the EMIT dataset (https://www.synapse.org/EMIT), which we hope will become a standard for evaluating new methods for multiplexed image analysis (similar to MNIST for classical machine learning). The dataset consists of 123 images spanning 34 different tissue types. Each image is approximately 3,000 x 3,000 pixels and 64 channels, with file size being ~1GB. |
| 10^5 - 10^6 cells, not sure about pixels, 10-100gb, 10-30 markers |
| TB |
| tens of thousands of cells per sample, 5-10GB per sample, whole transcriptome or 20 protein markers |

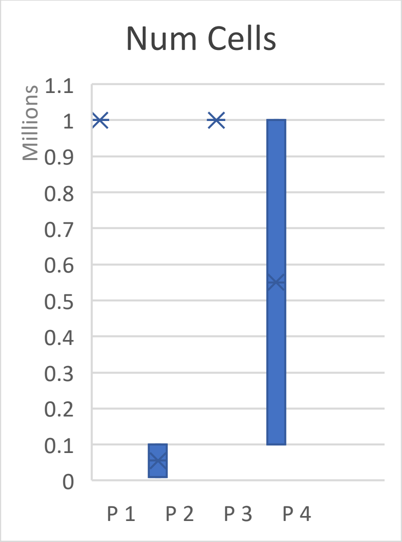


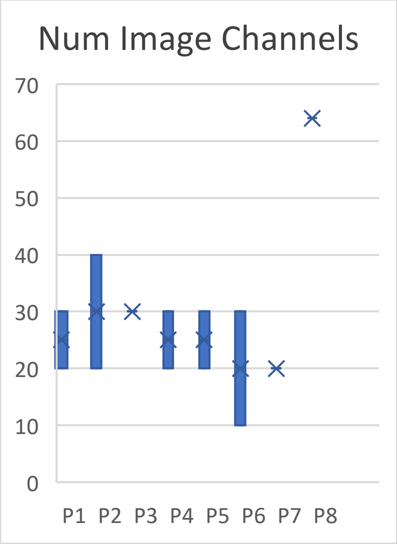


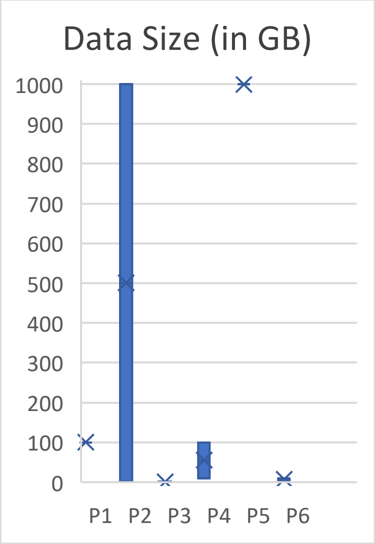


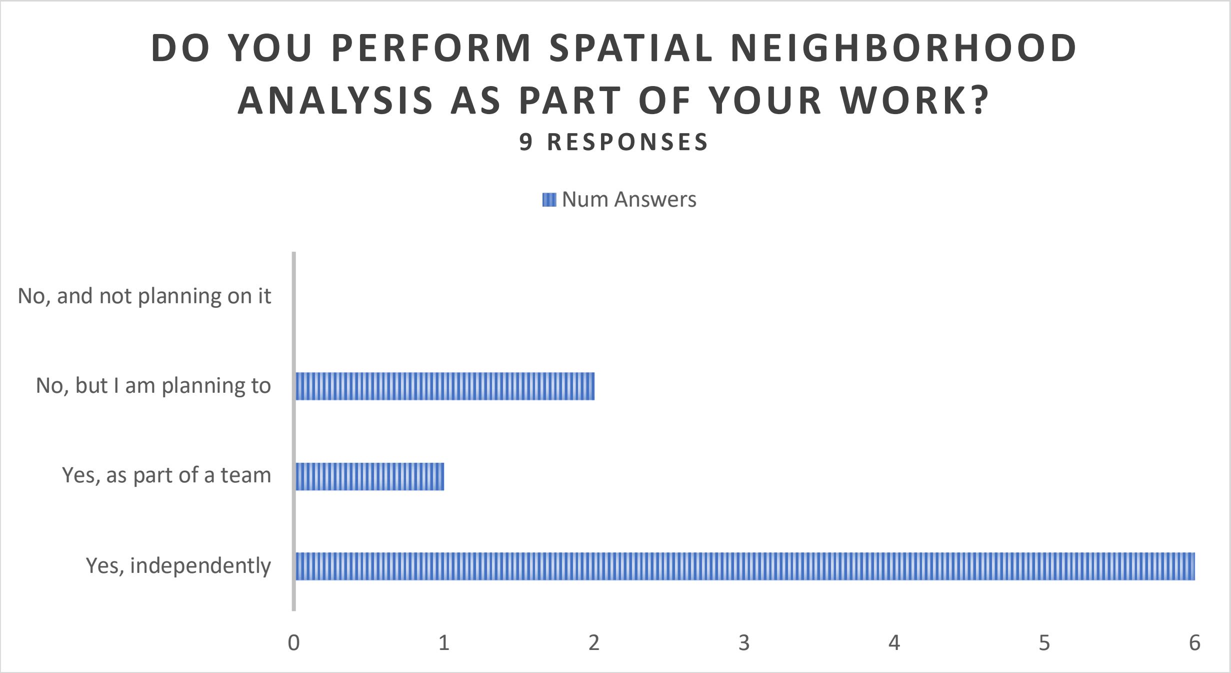


**What research/clinical questions are you asking when performing spatial neighborhood analysis?** **9 responses**

| |
|---|
| Which cell types attract/repel each other? |
| Which cell types tend to cluster in a network? |

| |
|---|
| Which ones are more independent? |
| What is the composition of different networks based on cell type? |
| Structure of the ecosystem (i.e., what cell types and cell states congregate) whether certain cell-types might be interacting (e.g. immune-tumor), or whether distinct regions of a tumor have different properties (e.g. composition, spatial patterns) |
| Network size/density? How does this change with treatment? |
| Difference in microenvironment between different tumors within same tissue? |
| How do neighborhoods change with disease sub-type? |
| How does cell phenotype differ between networks? |
| Level of tumor infiltration? |
| Tumor-TME interaction |
| Generally mapping immune populations or relating expression of proteins in tumor cells and surrounding cells. |
| Whether cell type can be inferred from morphological features and neighborhood information. The topic was part of the CSBC Image Analysis Hackathon in 2020. A manuscript outlining the findings is currently in preparation. |
| What spatial features correlate with clinical information |
| proximity of immune cells and cancer cells to infer putative interactions between tumor and immune cells |

**Do you perform spatial neighborhood analysis on a team, and, if so, who is on that team and what tasks to they perform (e.g. pathologist annotates images, computational biologists extract information from the image)?** 9 responses

| |
|---|
| Person X usually annotates tumors from H&Es or directly onto our CyCIF data. We integrate this info in our analysis and have been exploring different immune structures and how they relate to the tumor microenvironment |
| I am a part of .. a multi-disciplinary team that all tasks from image acquisition (experimental scientists) to image annotation (pathologists) to method development (computational scientists). |
| Yes, I am a pathologist and generally annotate regions, QC and refine staining, and analyze image results |
| Microscopists - helps in acquiring data |

| |
|---|
| Pathologists - help with interpreting spatial patterns |
| A pathologist annotates images, computational biologists extract information from the image and discusses with the team to refine the approach and interpret the data |
| There is no standard workflow |
| Biological question + Pathologists |
| Most of time, end-to-end analysis |
| Pathologist annotates image, biologist generates data, computational biologist extracts information from images. |

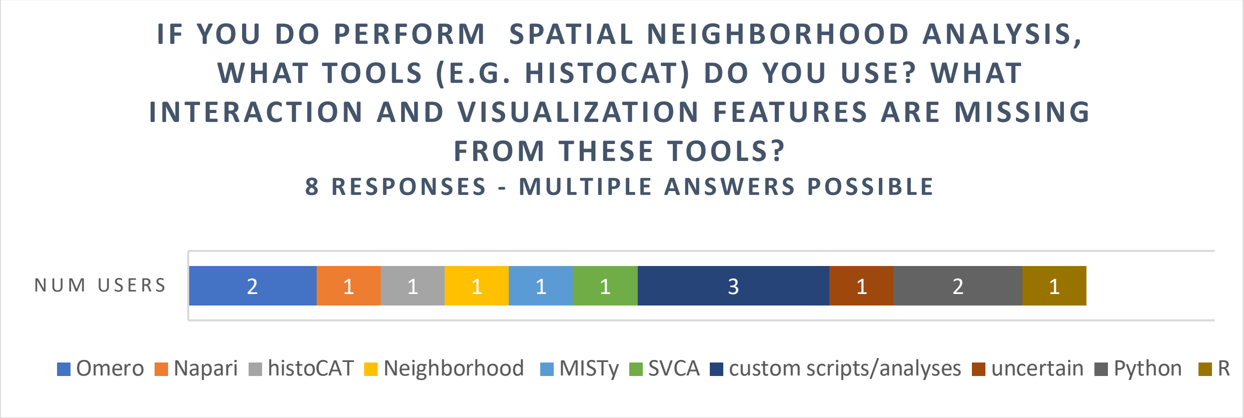


| |
|---|
| custom scripts. Limitation: is not being able to overlay neighbourhood clustering on images in real time. Not being able to visualize multiple images at the same time. |
| **MATLAB**. I'm satisfied with what I can code within MATLAB. |
| We use **Matlab**, **Omero**, and **Python** |
| **customized MATLAB scripts** |
| **R/Python** for method development. **Napari** for visualization of images. As a generic image viewer, napari doesn't understand file context. For example, it is not straightforward to apply a segmentation mask to the original image to define cell-specific regions. It is also not straightfoward to then annotate those regions with additional per-cell information (e.g., cell shape, size, etc.) or neighbor information (e.g., the distance between cells A and B is X). |
| Generally, these steps are done by collaborators, I'm not certain |
| **custom scripts**. Limitation: is not being able to overlay neighbourhood clustering on images in real time. Not being able to visualize multiple images at the same time. |

| |
|---|
| Members of the group perform **customized analyses**; spatial correlations are computed as the pearson correlation between a cell of group X and its kth nearest neighbor of group Y, for their respective variables x and y. Characteristic lengths are computed from exponential fit by least-square fitting (**MATLAB** in-built function lsqcurvefit). |
| MATLAB. I'm satisfied with what I can code within **MATLAB**. |
| e.g., **histoCAT**, **NeighboRhood** (https://github.com/BodenmillerGroup/neighbouRhood), **MISTy**, **SVCA** etc. |

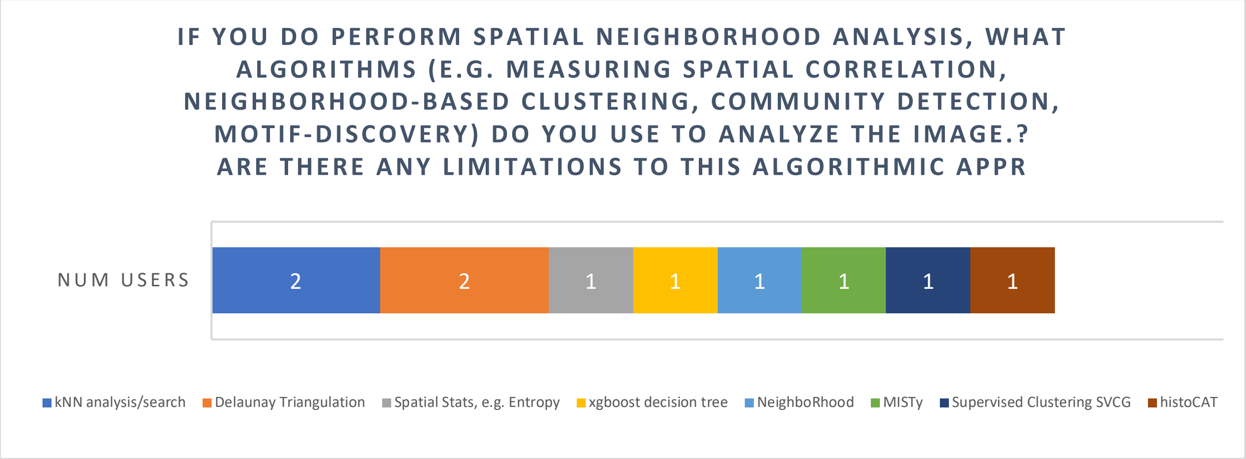


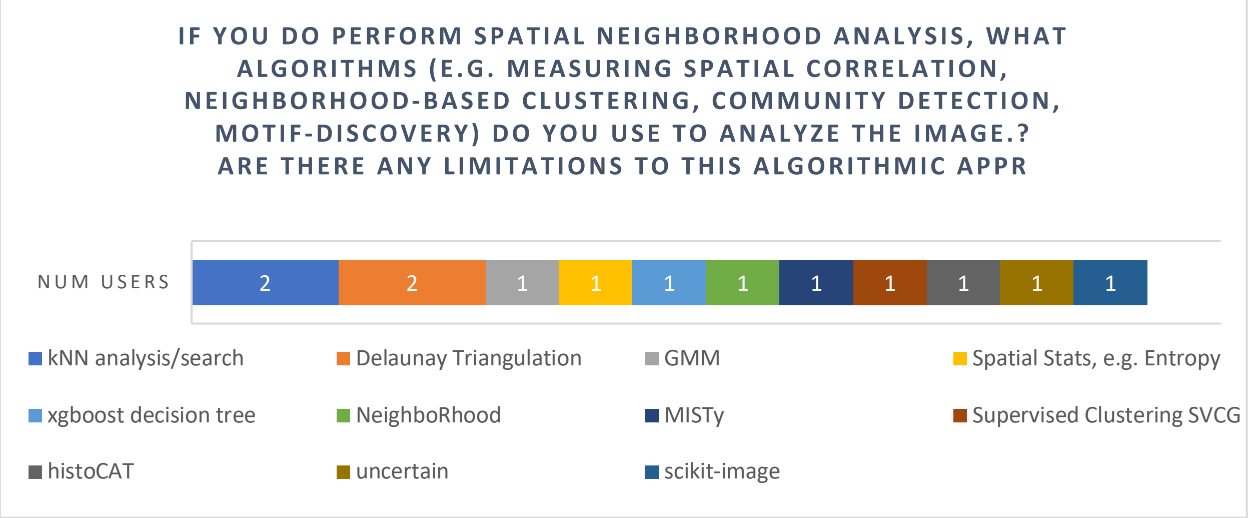


| |
|---|
| **kNN** searches, **Delaunay** triangulations, **Gaussian mixture modeling**, |
| **Leiden** Based Community Detection; Delaunay Triangulation; **nearest neighbor** analysis to look at cell to cell interactions regardless of communities |

| |
|---|
| Use most of (if not all) these approaches, also with other **spatial statistical** measures (eg. **entropy**). It really depends on the biological questions you try to address. |
| User friendly interactive tools would be very useful that permit on the fly assessments and immediate feedback that is mapped onto the image data. |
| For the question of cell type inference above, it was **xgboost** trained on morphological features (extracted via **scikit-image**) for the index cell and its closest five neighbors. |
| Also done by collaborators, not certain |
| e.g., **histoCAT**, **NeighboRhood** (https://github.com/BodenmillerGroup/neighbouRhood), **MISTy**, **SVCA** etc. |

**Briefly describe your current workflow from imaged data to insights. (e.g. image processing w/ mcmicro -> visual exploration -> cell type calling -> computing spatial statistics)** **9 responses**

| |
|---|
| mcmicro for image processing to retrieve single-cell information. Omero to look at images. Data analysis in python and use scatter plots and overlays on the images to interpret the data. |
| Highly dependent on project; various subsets of activities are all possible. |
| imaging--> ashlar--> Upload Omero--> Matlab--> Cropped Images for Ilastik Training --> Segmentation using probability maps from Ilastik --> single cell quantification in Matlab --> normalization of raw data through fitting it to Gaussian Mixture models --> cell typing --> network analysis |
| single-cell data extraction via McMicro or customized scripts --> binary gating --> cell type assignment (or clustering) --> spatial analysis |
| mcmicro or custom pipeline analysis -> visual exploration in omero -> cell type calling -> computing spatial statistic |
| mcmicro -> generating proximity graph based on (x,y) of individual cells -> defining feature vectors from morphological features of cells and their neighbors on the graph -> xgboost -> cross-validation -> examination of feature importance |
| Image processing with MCMicro and/or ASHLAR -> Upload to OMERO -> visual review on OMERO (or rarely ImageJ) by pathologist (me) -> marker QC and/or thresholding -> cell segmentation - > cell type calling by marker expression -> single cell analysis of marker expression (e.g. tSNE) - > spatial analysis (typically done by collaborator) - > data review and refinement. Often there are iterative cycles of single cell analysis and spatial analysis depending on emergent biological questions and insights. |
| MCMICRO -> Analysis with scripts -> Visualization in Napari/Omero<br><br>read alignment, mapping to spatial barcodes, dimensionality reduction, clustering, cell type mapping (RCTD, Seurat) using single cell reference dataset |

##### Are there any standards or best practices in your field to visualize spatial neighborhood information (conventional visual encodings, charts, and graphs)? 7 responses

| |
|---|
| No |
| Honestly, I am pretty simple and really enjoy a simple scatter plot where you plot individual cell types in different colors preserving their spatial relationships to get a quick and dirty idea of what the neighborhoods look like; Definitely open to more sophisticated ways of looking at the data |
| It's a topic of ongoing research, and best practices are yet to be established. |
| This is probably better answered by those directly generating the analyses. So far, my projects have typically employed length scale analysis, nearest neighbor analysis (e.g. kNN), Delaunay clustering, etc. |
| no. I have generally seen cell-cell interactions displayed as a heatmap or cirus plot. |
| these are still developing |
| none, but could use more from other disciplinaries (eg. GIS). |

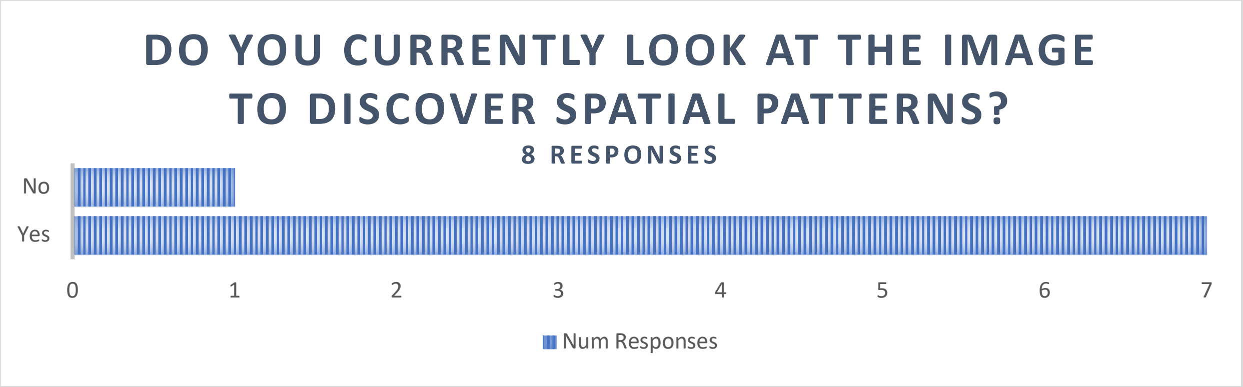


One participant added an additional comment:

| |
|---|
| Yes, this is critically important. I am a pathologist so this is often one of my main responsibilities. In my experience on multiple projects qualitative visual review of imaging data is critically important to identify erroneous staining, threshold markers for cell type calling, identify subjective trends for quantitative analysis, and re-check data-driven conclusions. This is often overlooked leading to spurious conclusions, incorrect cell calling, and thereby incorrect neighborhood analyses. Analysis of tissue specimens is simultaneously deceptively straightforward but actually highly complex, and marker expression and distribution is not easily analyzed without input from pathologists who have seen numerous variations in marker and cell patterns within anatomic context in practice. I would be highly cautious about interpreting tissue specimens without pathology input, there are many pitfalls including reative atypia, known off-target staining, unusual or subtle tumor phenotype, etc. that are not easily appreciated without extensive training. |

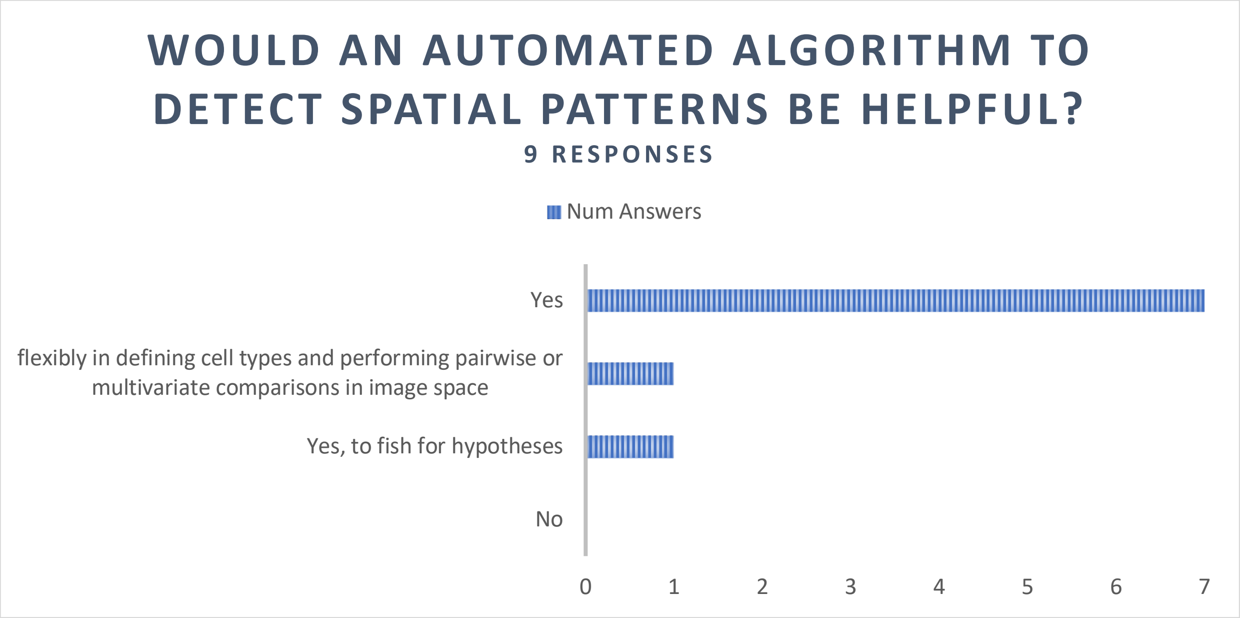


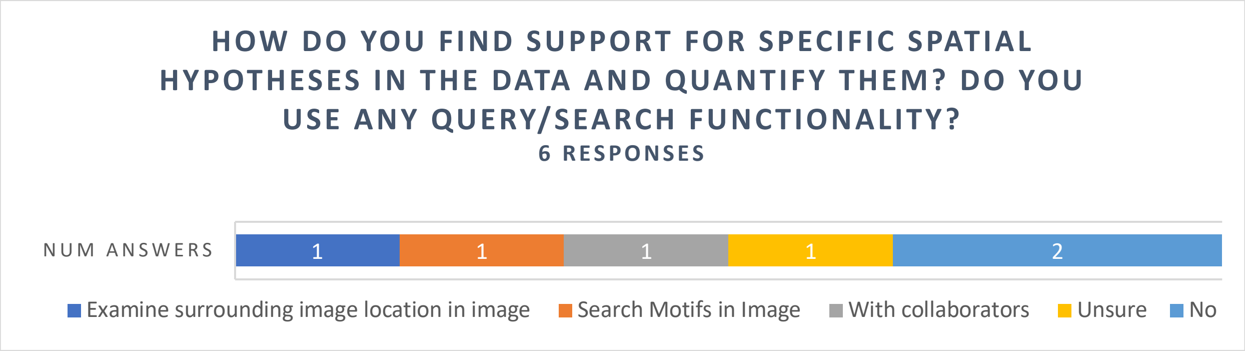


| |
|---|
| For the question of cell type calling, we identify outliers (i.e., with high/low prediction confidence) and examine the corresponding location in the images, in an attempt to understand why the cells were classified correctly or incorrectly. |
| I have been fortunate to be extensively involved with the development and implementation of CyCIF and spatial biology throughout my training at HMS, and so there are numerous collaborators who have been generously supportive with providing insight and guidance with all aspects of image analysis on my projects and theirs. |
| currently no but as explained previously, there are certain hypothesis we would like to test and visualize those interactions on the image. |
| Search of reference motifs both within as well as across specimens will be needed what |
| Not sure about this |
| Not at that stage in the analysis yet |

**Would the ability to draw a region of interest in the image and find similar spatial regions support your work? Can you give a use-case example? 9 Responses**

| |
|---|
| I think this would definitely be very interesting. In one of my current datasets, we are very interested in the B cell niches that we are seeing in certain lung cases. They can be centers for t cell activation and are associated with better outcomes. It would be very cool to easily stratify these different B cell niches into separate categories depending on the different markers expressed. Maybe some are more dysfunctional than others. Maybe some are closer to exhausted t cells than others. |
| Yes, if there is a specific region that contains many misclassified cells (e.g., because of an artifact in the data), it would be help to know if similar artifact are present elsewhere. |
| Yes, this would be extremely useful. Actually, I am one of the main people drawing regions of interest for regional and spatial analysis as a pathologist. I would be happy to discuss further outside of this survey. For instance, I have defined specific tumor regions by differentiation state or anatomic invasion correlating with clinical parameters, defined specific regions of interest for immune profiling, or specific regions of interest within normal or tumor regions. We have leveraged this to study unique regions within tumors (e.g. defining stem-like whorled regions in craniopharyngioma), examined and compared the spatial profile and distribution of immune populations within tumor stroma, at the tumor-stromal interface, and intratumorally, done global analysis of immune populations in tumors, examined tumor and immune marker distribution across defined ROIs. It is very important to be able to define complex ROIs within tissue specimens (i.e. not just squares or circles, but free-hand ROIs), and extract those regions for analysis. Tissue structures are highly complex and require complex ROIs for precise analysis. Some phenotypes will not be able to be analyzed appropriately with simple circle/square ROIs. There are innumerable examples from pathology and tumor biology that could be discussed. |
| yes (this seem to be a repetitive question) |
| Yes - identification of community of cells (cells in two cell states) across tumors and mapping them to general location (border, etc.) will be helpful |
| Yes, although the definition or existence of "similar" spatial regions is highly subjective. |
| Would be awesome for e.g., immune infiltration or unique features at tumor margins etc. |
| yes, could be useful to identify ROIs (eg tumor versus stroma) or even different cell types |
| Yes |

**Would the ability to query for specific cell to cell interactions (e.g., Tumor-CD8T cell interaction) and quantify/visualize the results support your work? Can you give a use-case example?** 9 Responses

| |
|---|
| Absolutely. It would be especially helpful to look at what kinds of tumor cells are closer to CD8 t cells. Are they PD-L1+ and therefore potentially inducing dysfunctional programs? Are they proliferative (i.e. PCNA+ or Ki-67+)? |

| |
|---|
| Are the tumor cells in dense networks? What is the tumor grade? Are the CD8 t cells dispersed? Clustered in certain regions? What are the phenotypes of CD8 t cells? |
| Yes. This is in fact the main question of the cell type prediction task: is it possible to recognize tumor-infiltrating immune cells based on the shape of those cells and the fact that they are surrounded by tumor cells? |
| Yes, this would be extremely useful. The interaction of tumor and CD8 cells as you've noted is a particularly important example, as interaction of tumor cells and cytotoxic T cells (CD8+; the main cell type responsible for immune-mediated tumor killing) is of special interest in tumor biology and pathology. Another similar example would be tumor-NK cell interaction. The interaction of PD-1+ immune populations and PD-L1+ populations (tumor and immune) to examine checkpoint signaling. The interaction of immune populations and DNA damage or innate immune markers. The interaction of signaling pathway markers in different cell populations (see our craniopharyngioma paper with regard to signaling pathways and immune signaling in whorled cells). Interaction of stem-like populations and other cells in both neoplastic and non-neoplastic settings. There are innumerable examples that could be named. |
| certainly. by visual inspection some patterns are very clear but sometimes they are not and being able to identify those more encrypted patterns could be useful. |
| yes. |
| Yes. this would be helpful. |
| Absolutely! |
| Yes, e.g. PD-1 & PD-L1 interaction |
| Yes, absolutely |

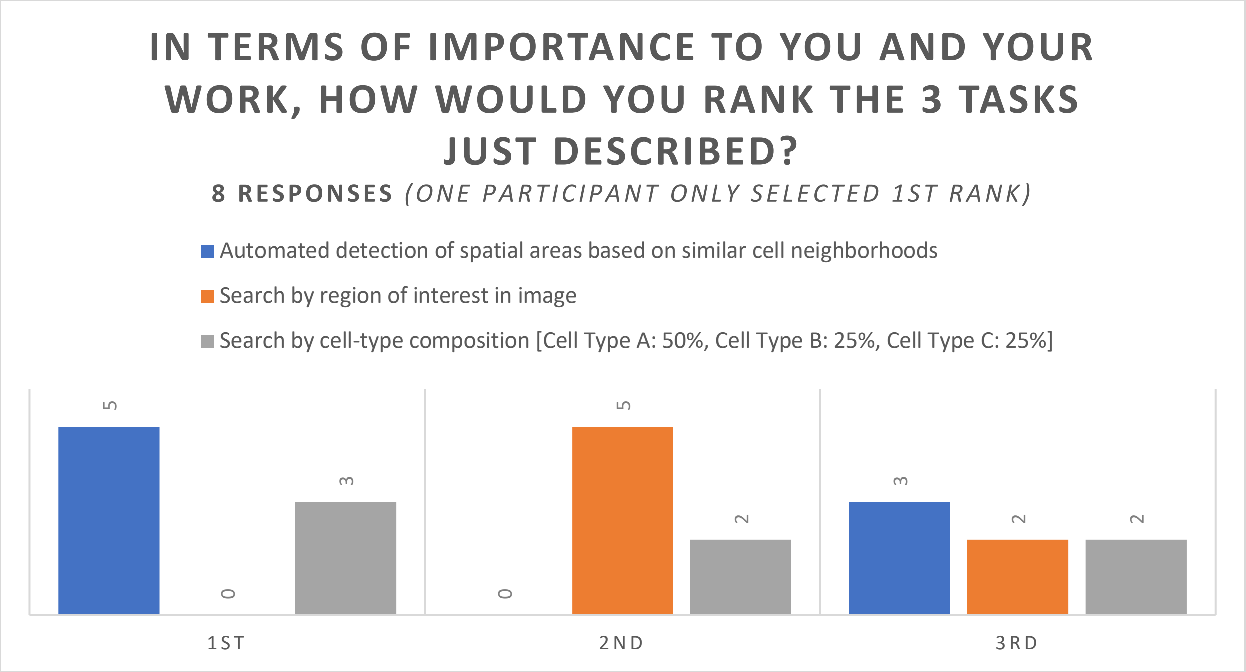

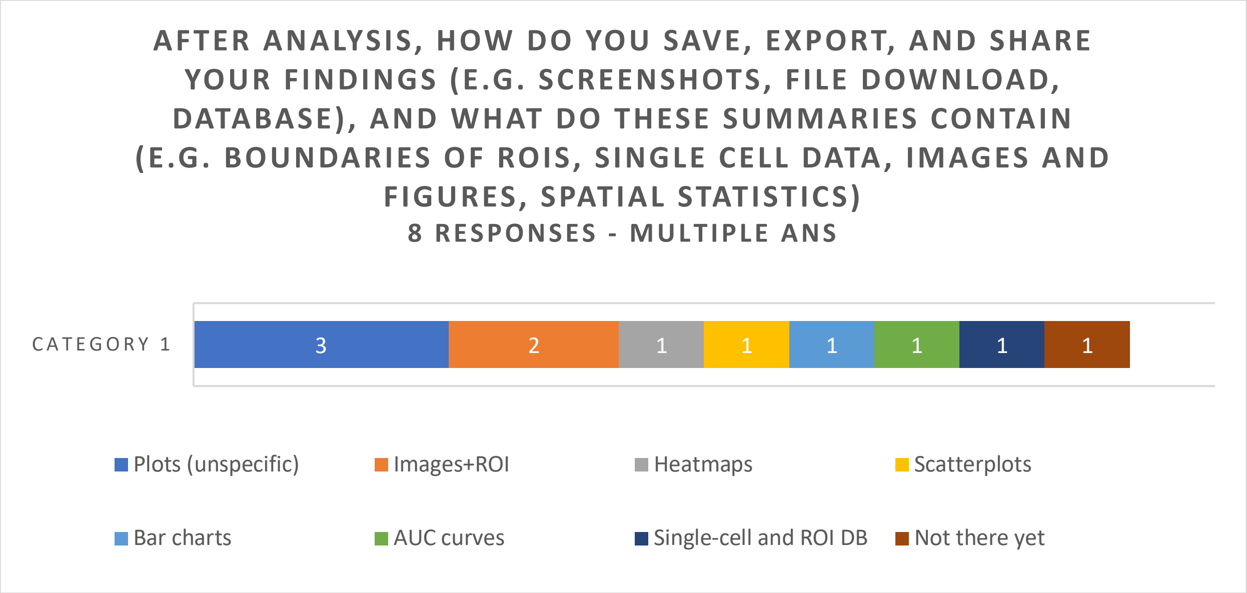


| |
|---|
| Omero **ROIs**, Matlab **scatter plots** of different cell types, k means clustering **heat maps**, **bar charts** with cell type info, bar charts showing median distance between cell types, distance distributions highlighting how close certain cells are to each other, images from matlab, etc. |
| Programmatically-generated plots (e.g., AUC curves) are exported to .pdf, while numeric data (e.g., predicted cell types) are exported to .csv. |
| Yes, exporting images and ROIs, exporting image screenshots, single cell marker databases (globally and ROI based), spatial statistics |
| This is done by plots of spatial correlation and length scales coupled to high resolution images to display the features. |
| Do you mean figures in papers for "summaries", or do you mean data depositories (which are not succinct summaries)? |
| Usually just the figures and the corresponding scripts |
| figures or table generated via customized scripts |
| Not there yet |

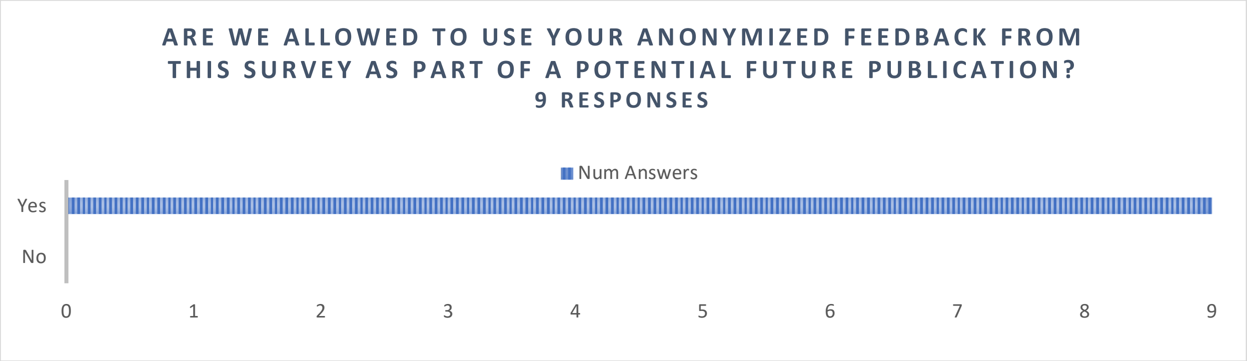

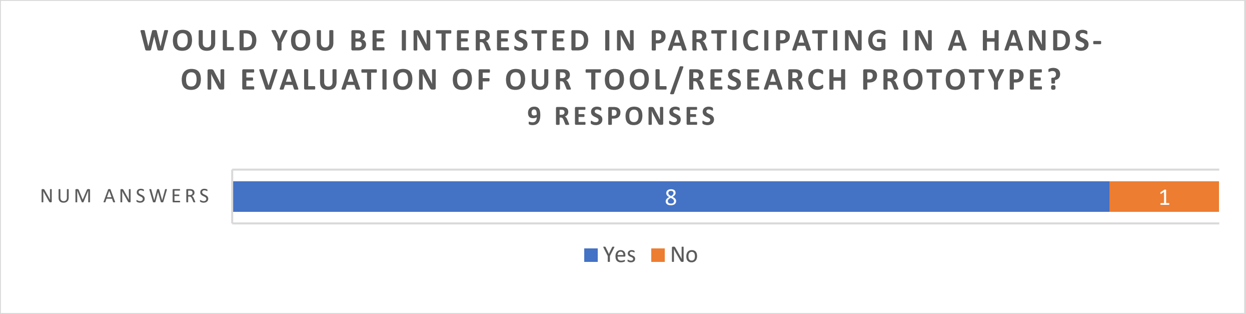


##### Is there anything else you think would be helpful to share? 2 responses

| |
|---|
| Yes, happy to discuss elsewhere anytime as well. |
| Some motifs differ in size from small to increasing large and it would help to map molecular gradients across motifs of varying size |
